# Epiblast morphogenesis is controlled by selective mRNA decay triggered by LIN28A relocation

**DOI:** 10.1101/2021.03.15.433780

**Authors:** Miha Modic, Igor Ruiz de Los Mozos, Sebastian Steinhauser, Emiel van Genderen, Silvia Schirge, Valter Bergant, Joel Ryan, Christopher B Mulholland, Rupert Faraway, Flora C Y Lee, Tajda Klobučar, Juliane Merl-Pham, Stephanie M Hauck, Micha Drukker, Sebastian Bultmann, Heinrich Leonhardt, Heiko Lickert, Nicholas M Luscombe, Derk ten Berge, Jernej Ule

## Abstract

The embryonic progression from naïve to primed pluripotency is accompanied by the rapid decay of pluripotency-associated mRNAs and a concomitant radical morphogenetic sequence of epiblast polarization, rosette formation and lumenogenesis. The mechanisms triggering and linking these events remain poorly understood. Guided by machine learning and metabolic RNA sequencing, we identified RNA binding proteins (RBPs), especially LIN28A, as primary mRNA decay factors. Using mRNA-RBP interactome capture, we revealed a dramatic increase in LIN28A mRNA binding during the naïve-rosette-primed pluripotency transition, driven by its nucleolar-to-cytoplasmic translocation. Cytoplasmic LIN28A binds to 3’UTRs of pluripotency-associated mRNAs to directly stimulate their decay and drive lumenogenesis. Accordingly, forced nuclear retention of LIN28A impeded lumenogenesis, impaired gastrulation, and caused an unforeseen embryonic multiplication. Selective mRNA decay, driven by nucleo-cytoplasmic RBP translocation, therefore acts as an intrinsic mechanism linking cell identity switches to the control of embryonic growth and morphogenesis.

---

Pluripotent cells progress through several rapid developmental intermediates before acquiring trilineage differentiation potential. Over a span of 24 hours surrounding embryo implantation, major morphological transitions take place: the naïve pluripotent epiblast cells (mouse embryonic day E4.5) polarize and arrange into a rosette, then irreversibly transition into primed pluripotency while the rosette undergoes lumenogenesis to generate the egg cylinder and pro-amniotic cavity (E5.5) (1–3). Rapid dismantling of the naïve pluripotency gene network in a brief time window (E4.9–5.1) is required to progress through the rosette stage and initiate lumen formation (1). Failure to do so interferes with lumenogenesis in mouse and human embryos (1, 4).

The mechanism that triggers and coordinates this cascade of dramatic molecular, cell-fate and morphological transitions of early embryonic development remains poorly understood. We hypothesized that enhanced mRNA decay is crucial for the rapid drop in mRNA abundance of naïve pluripotency genes that has been observed during naïve-to-primed transition (5). To define the critical period when naïve gene expression is downregulated, the naïve-to-primed transition was recapitulated by transferring naïve embryonic stem cells (ESCs) to medium containing FGF2 and ActivinA (6). After 12 hours, a rapid and coordinated depletion of naive pluripotency mRNAs was observed (7), followed by a drop in the abundance of the corresponding proteins (Fig. 1A, S1A-B). To determine whether this depletion resulted from selective mRNA decay, we employed metabolic RNA sequencing (SLAMSeq) (8) to directly measure half-life of transcripts in naïve ESCs and upon 12hours of naive-to-primed transition (priming ESCs). This revealed that transcripts encoding genes downregulated in primed pluripotency (developmental genes) become significantly less stable in priming ESCs (Fig. 1C).

**Figure 1.**
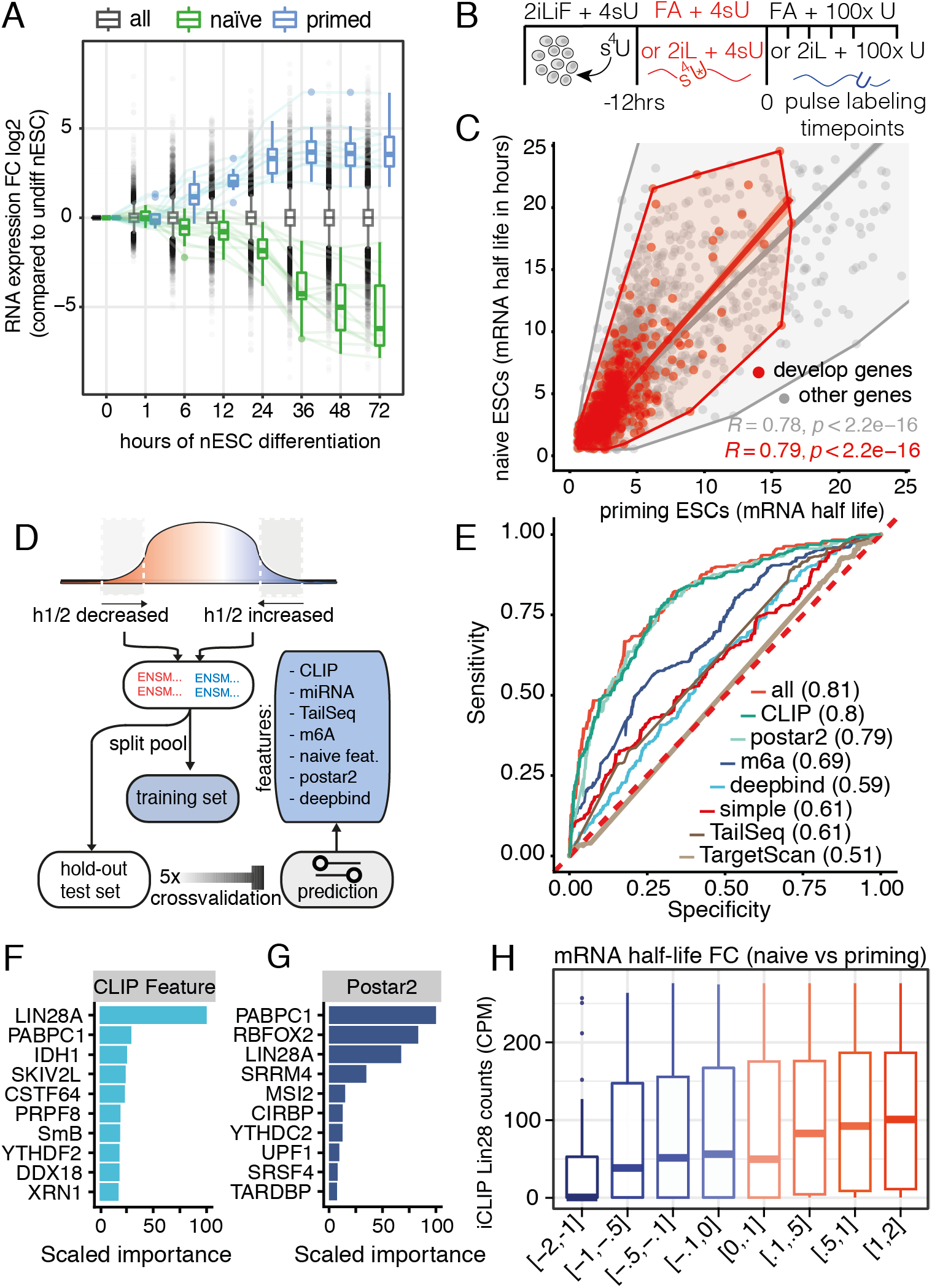
Machine learning predicts LIN28A-dependent control of developmental switch in mRNA stability. (A) Temporal dynamics of relative mRNA levels (compared to 2iLIF at 0hrs) of naïve (green) and primed (blue) genes downregulated during hours of ESC naïve-to-primed differentiation (7). Error bars represent SEM. (B) Protocol for SLAMseq measurements of mRNA half-life during naïve-to-primed pluripotency transition (C) mRNA half-life of a 6,052 genes (R2 > 0.6) determined at high accuracy at naïve-to-primed turning point (x-axis) and naïve state (y-axis). Transcripts that are downregulated upon 48 hours of naïve-to-primed transition are labeled red (developmental genes as determined in (7)). Colored lines represent the Pearson correlation. (D) Machine learning framework for classifier implementation using the top 1,000 most UP/DN regulated genes are randomly split into a training set and a hold-out test set. Gradient boosted trees are trained on the features of the training set using 5 times repeated 5 fold cross-validation while performing hyperparameter optimisation using grid search. (E) ROC curve performance benchmark of all resulting binary classifiers trained on different feature sets. The area under the ROC curve (auROC) is indicated for each model in the legend. (F-G) Feature importance ranking, based on the relative influence, for the two best performing classifiers based on (F) mESC CLIP (n=62) and (G) Postar2 CLIP-seq database (10). (H) Normalized LIN28A binding (y-axis) to 3’UTRs of transcripts against their relative rank order (x axis) according to change in their mRNA halflife during naïve-to-primed transition..

To uncover features that predict the changes in mRNA halflife between naïve and priming ESCs, we trained machine learning classifiers based on gradient boosted trees using various feature sets including mRNA dinucleotide frequencies, published and new datasets on RNA binding sites of RBPs determined with crosslinking and immunoprecipitation (CLIP), miRNA target sites, m6A sites, TAIL-seq (9), Postar2 features (10), and DeepBind (11) predictions of RBP binding within 3’UTR (Fig. 1E). The classifiers were trained to distinguish decreased (DN) and stable/increased (UP) half-lives using the top 1,000 DN/UP transcripts. We identified RBP binding as the most predictive feature of the change in mRNA stability upon naive-to-primed transition (Fig. 1E). Surprisingly, no association was found with specific miRNA families, AU-rich elements, polyA-tail length or other features (Fig. S2A). Classifiers trained on the enrichment of RBP binding to 3’UTRs were able to correctly predict the change in half-life of mRNAs previously unseen by the model (auROC=0.8, accuracy=0.72, Matthews correlation coefficient (MCC)=0.43) (Fig. 1F, Fig. S2B-C). Feature importance analysis, based on the relative influence, indicated LIN28A and PABPC1 as top-ranking RBPs to predict observed half-life changes (Fig. 1G-H). These rankings were additionally verified by a permutation based feature importance (Fig. S3). Accordingly, the extent of LIN28A binding to mRNAs in ESCs is proportional to their decreased half-life (Fig. 1H).

Given that RBP binding to 3’UTRs is the primary feature predicting developmental mRNA stability, we performed global mRNA-RBP interactome capture (12, 13) in parallel with whole-cell proteomics to identify RBPs that dramatically change in mRNA binding during early development (Fig. 2A). We hypothesised that a subset of RBPs acting as key developmental regulators will increase mRNA binding during the naïve-to-primed transition. For this purpose, we benchmarked and generated a homogeneous population of naïve ESCs, rosette stem cells (RSCs) (4), primed Wnt-inhibited epiblast stem cells (EpiSC) (14) and the earliest lineage-specified mesodermal progenitors (15), FACS-sorted using the T-eGFP marker. We adapted mRNA-RBP interactome capture (16) to make it applicable to FACS-sorted progenitors with a robust RBP discovery rate (Fig. 2G, S4). We validated the specificity of the mRNA-RBP interactome enriched over non-cross-linked and naked beads controls with GO terms (Fig. S5A-D). Significantly, 9, 14 and 48 RBPs exhibited changes specifically at the level of mRNA binding in RSCs, EpiSCs or mesodermal progenitors compared to their binding in naïve ESCs, respectively (Fig. S5E-G). These are considered dynamic RBPs as their enrichment could not be accounted for by the changes in total protein abundance (absolute lasso-based log2 discrepancy > 0.25 with associated p value < 0.001, for more information see Methods). Among these, LIN28A exhibited the highest relative increase in mRNA binding in RSCs, EpiSC, and mesodermal progenitors, which validated and extended our findings from the machine learning classifiers (Fig. 2B S5E-G). We hypothesised that the change in LIN28A mRNA binding could be driven by relocalisation of the protein during the naïve-to-primed transition. Therefore, we tagged the endogenous Lin28a gene with eGFP and tracked the protein with live cell imaging (Fig. S6A-B). As reported for the inner cell mass of early blastocyst (17), we observed LIN28A-GFP predominantly localized in the nucleolus of naïve ESCs. However, Lin28A translocates to the cytoplasm upon the onset of embryonic priming (Fig. 2C-D). This translocation was confirmed by immunofluorescence imaging and by western blot analysis of nuclear and cytoplasmic fractions (Fig. 2E, S6C-D). The increase in relative mRNA binding of LIN28A during the naïve-to-primed transition was further corroborated by RNA-protein crosslinking and immunoprecipitation (iCLIP) (Fig. S6E). The nucleolar-to-cytoplasmic translocation and increased mRNA binding strongly implicate LIN28A in the selective decay of pluripotency mRNAs early during the naïve-to-primed transition.

**Figure 2.**
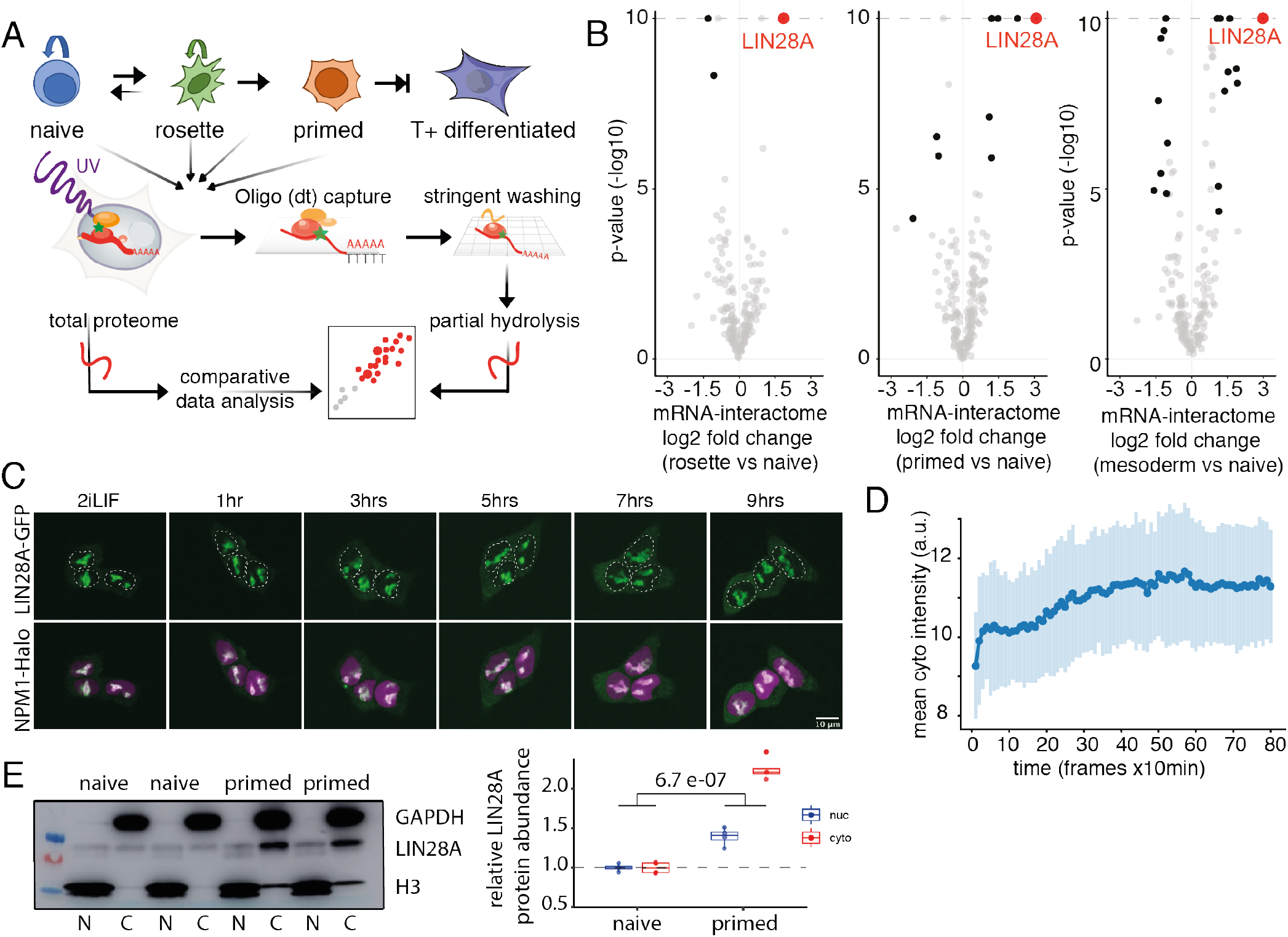
Figure 2: LIN28A translocates to cytoplasm and increases mRNA binding upon naïve-to-primed transition. (A) The workflow of the mRNA-RBP occupancy assay (B) Volcano plots depicting the difference in RBPome of bona-fide RBPs (thresholding determined in Fig. S5) between rosette stem cells and ESCs (left), EpiSCs and ESCs (middle) and mesodermal progenitors and ESCs (right panel). RBPs marked black fulfilled: absolute log2 FC > 1 and p < 0.0001 using lasso-based linear model. (C-D) Localization dynamics of endogenous LIN28A-GFP in response to embryonic priming during confocal time-lapse imaging over 12 h (10 min intervals) and quantification of mean cytoplasmic intensity of the endogenous LIN28A-GFP signal. Scale bar: 10 μm. (E) Quantification with the results of four independent biological replicates displayed of Cytoplasmic, “C”, and nuclear, “N”, fractions from ESCs and EpiSCs analyzed by immunoblot detection using the indicated antibodies. A significant interaction term was identified by 2-way ANOVA, confirming that the nuclear-cytoplasmic localisation of Lin28a changes across the naïve-to-primed transition.

To directly assess the role of LIN28A in regulating mRNA stability, we primed LIN28A knockout (KO) and WT ESCs for 12hrs and conducted SLAMseq metabolic labeling to measure mRNA stability of 6,276 transcripts (R2 > 0.6) (Fig. S7A-B). Importantly, the decrease in mRNA stability observed after 12 hrs in primed conditions did not occur in LIN28A KO cells (Fig. 3A), which was most apparent for mRNAs that normally display a strong decline during the naïve-to-primed transition (defined in Fig. S1A) (Fig. S7C-D). Moreover, the changes in transcript stability after 12 hrs in primed conditions correlated highly with the changes between WT and LIN28A KO priming ESCs, indicating that the developmental decrease in mRNA stability is indeed driven by the increased abundance of cytoplasmic LIN28A (Fig. S7E-F).

**Figure 3.**
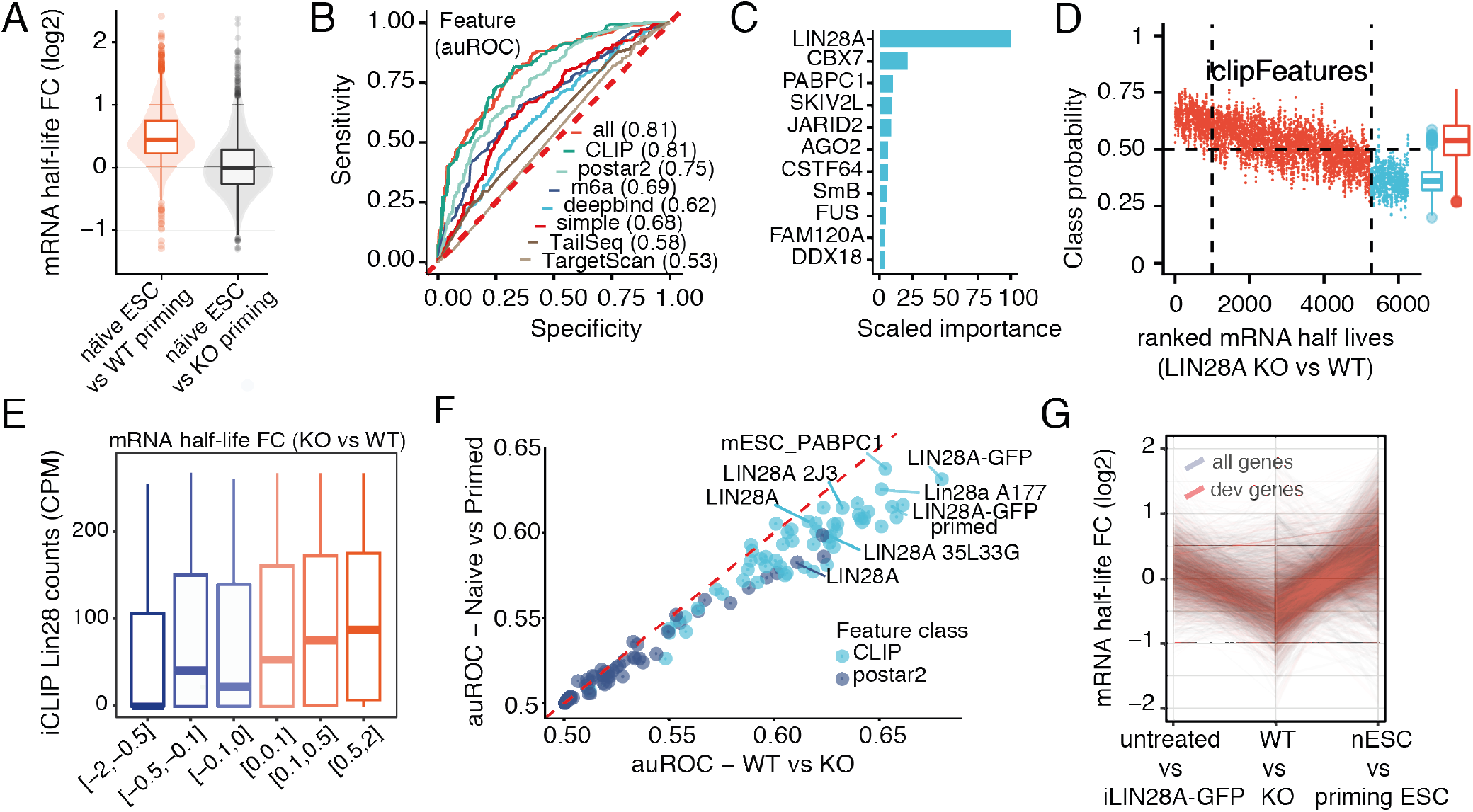
LIN28A binding decreases the half-life of its mRNA targets. (A) Violin-plots depicting mRNA half-life fold change upon priming in WT and LIN28A KO ESCs compared to naïve ESCs. (B) ROC curve performance benchmark of all resulting binary classifiers (WT vs LIN28A KO) trained on previously introduced features (Fig. 1). The area under the ROC curve (auROC) is indicated for each model in the legend. (C) Feature importance ranking, based on the relative influence, for the best performing classifiers based on mESC CLIP combined dataset. (D) Ranking of all half-life changing transcripts from UP towards DOWN upon LIN28A KO compared with the class probability predicted with the binary model trained using CLIP features. 1000 most destabilized transcripts are labeled blue. Class probabilities were smoothed using the running mean (k=10). (E) Normalized LIN28A binding (y-axis) to 3’UTRs of transcripts against their relative rank order (x axis) according to change in their mRNA half-life upon LIN28A KO. (F) Comparison between auROC statistics computed for each RBP CLIP to distinguish either transcript stability changes in LIN28A KO vs WT (x-axis) or naïve vs prime (y-axis). (G) Line-plot depicting the half-life fold change upon LIN28A KO, iLIN28AGFP transient overexpression and in naïve ESCs compared to priming ESCs of mRNAs downregulated upon 48 hours of naive-to-primed transition (red) or all mRNAs (grey).

We then validated our machine learning framework by showing that LIN28A 3’UTR binding is the most important feature predicting mRNA half-life changes between LIN28A WT and KO cells (Fig. 3B-C, S7G-H, S8A). Furthermore, we observed a linear relationship between the CLIP-based predictions of mRNA classes and the differential mRNA stability comparing WT and LIN28A KO cells (Fig. 3D, Fig. S8B). Additionally, we observed a quantitative relationship between LIN28A binding and the extent of change in mRNA stability (Fig. 3E). To compare the feature importance for transcript half-lifes between changes in LIN28A KO/WT priming ESCs and the naïve/priming WT ESCs, we used the 3’UTR CLIP scores for each RBP to distinguish respective conditions resulting in an auROC for predicting mRNA halflife. This showed overall a high correlation of RBP feature importance between the aforementioned two sets of half-life changes, with the various published LIN28A iCLIP datasets interchangeably performing best in both cases (auROC>0.60, Fig. 3F). Thus, half-life changes can be predicted by LIN28A binding to 3’UTRs.

To exclude the possibility that changes in stability are a secondary result of LIN28A-caused cell fate changes, we generated stable doxycycline-inducible iLIN28A-GFP ESCs with pronounced cytoplasmic LIN28A accumulation to assess the impact of acute 10 hr LIN28A induction on mRNA stability in naïve ESCs (iLIN28A-GFP, Fig. S9A-D). Notably, this decreased the half-life of LIN28A-bound mRNAs in the same direction as 12 hrs in primed conditions (Fig. S9E). The LIN28A-GFP induced destabilization was pronounced for transcripts downregulated during the naive-to-primed transition (defined in Fig. S1A) and, as expected, was reciprocal to the transcripts that gained stability in LIN28A KO cells (Fig. 3G). To exclude the indirect contribution of miRNA-dependent stability effects (18, 19), we observed that mRNA destabilization preceded expression of the let7 miRNA family, thus it could not be mediated indirectly by changes in let-7 biogenesis (Fig. S9F). This agrees with reported LIN28A-induced gene expression changes prior to any effect on let-7 biogenesis (20). Combined, these findings argue that cytoplasmic localization of LIN28A is sufficient to drive pluripotency mRNA destabilization in a let7-independent manner.

Next, we investigated whether the LIN28A-mediated changes in mRNA stability are responsible for the dissolution of the naïve pluripotency gene network. First, we used CRISPR-Cas9 editing to excise a cluster of LIN28A binding sites from the 3’UTR of Sox2 mRNA. Sox2 is abundantly expressed in the naïve state, but its RNA and protein levels decline upon the transition to primed pluripotency (Fig. S1B, S4D) (7). Loss of the LIN28A binding cluster significantly increased the abundance of SOX2 protein upon naïve-to-primed transition (Fig. 4A-B). Second, we used iLIN28A-GFP ESCs and analyzed transcriptome changes upon 6 hours of LIN28A-GFP induction while maintained in naïve conditions. Strikingly, naïve pluripotency mRNAs were specifically downregulated (Fig. 4C), with concomitant decrease in the protein levels of naïve pluripotency factors (Fig. 4D). Additionally, rapid induction of cytoplasmic LIN28A in naïve conditions is sufficient to deplete the majority of transcripts that are downregulated during naïve-to-primed transition (Fig. S9G). Acute induction of cytoplasmic LIN28A-GFP induced the primed marker SSEA4 in naïve pluripotency conditions. Conversely, it reduced the naïve marker SSEA1 (Fig. 4E). In conclusion, rapid nuclear-to-cytoplasmic translocation of LIN28A promotes degradation of its target pluripotency-associated mRNAs to facilitate the naïve-to-primed transition.

**Figure 4.**
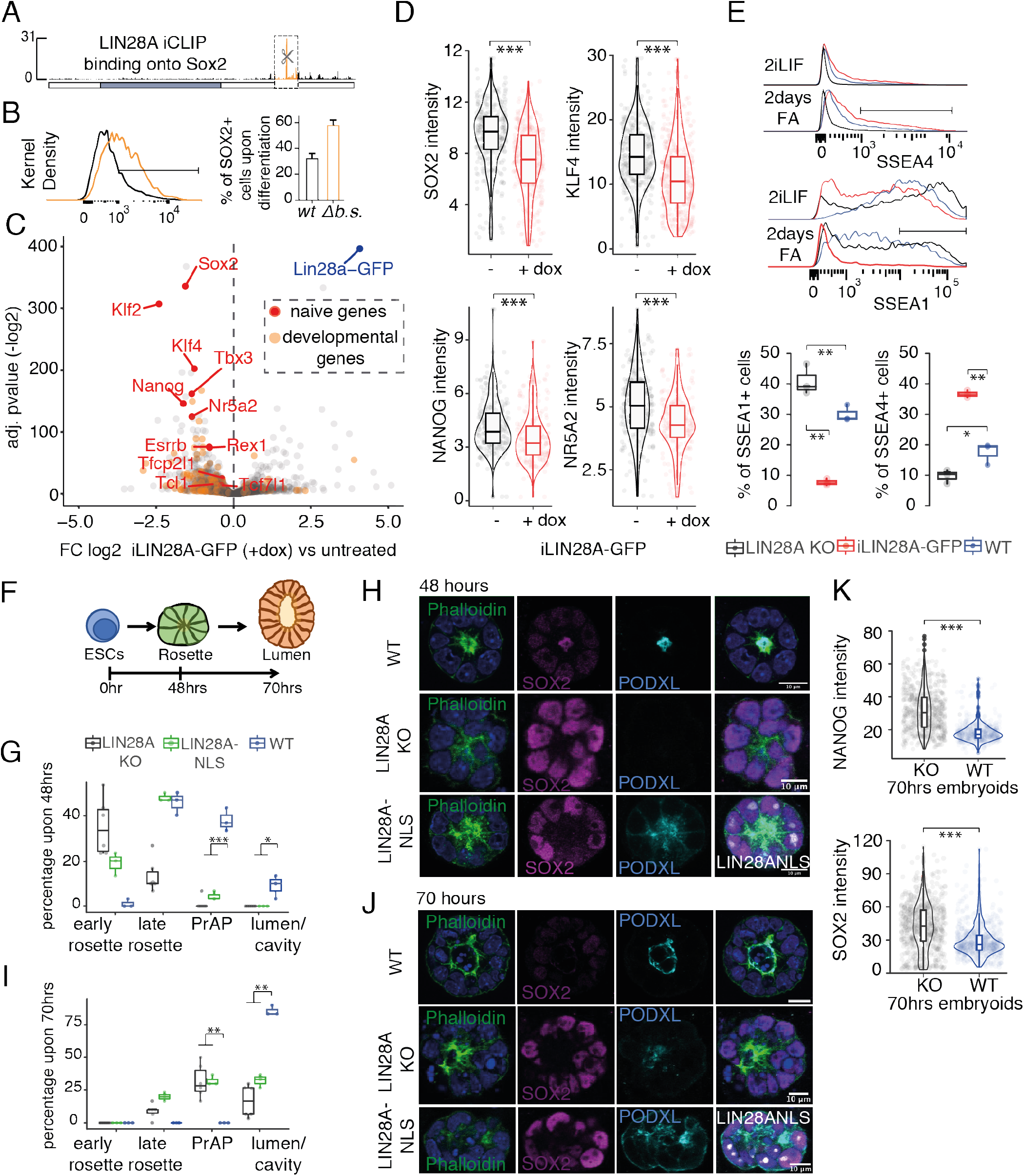
LIN28A translocation enables naïve-to-primed transition and facilitates lumenogenesis. (A) An illustration of the Sox2 transcript displaying a cluster of CRISPR-Cas9 excised LIN28A binding sites (Sox2Δb.s.) (B) Representative intracellular SOX2 flow cytometry and quantification of primed WT and Sox2Δb.s mESCs that lack the LIN28A binding cluster (n=3). (C) Volcano plot displaying gene expression fold changes and their respective statistical score (BH-adjusted p value, Fisher’s exact test), comparing tdoxycyclin-induced iLIN28AGFP and untreated undifferentiated mESCs (n = 4 biological replicates per condition, two clones). Naïve pluripotency markers are labeled in red (marked factors fulfilled p < 210) and genes downregulated during naïve-to-primed conversion (defined in Fig. S1A) are labeled in orange. (D) Fluorescence arbitrary units of immunostained naïve markers SOX2, KLF4, NANOG, NR5A2 spheres. n = >200. Two-sided t-test; ***P = <0.0001 (E) Representative flow cytometry analyses of naïve (2iLif) and primed (2 days FA) ESCs that were treated with doxycycline (iLIN28A-GFP), left untreated (LIN28A KO) and compared to WT and immunostained for SSEA-1 and SSEA-4 The mean of three independent primed experiments is depicted below (error bars, SD); Two-sided t test, ***p < 0.001, *<0.05 (F) Sequence of the morphogenetic events that drive the peri-implantation development (G-H) Rosettes and in vitro lumens 48hrs after seeding WT, LIN28A KO and LIN28A-GFP-NLS ESCs in BME in primed conditions, stained for indicated markers and (I) the percentage of ESCs generating rosettes and lumens 48hrs after seeding. Remaining aggregates were unpolarized, combined amounting to 100(I-J) Rosettes and in vitro lumens 70hrs after seeding WT, LIN28A KO and LIN28A-GFP-NLS ESCs in BME in primed conditions, stained for indicated markers and (I) the percentage of ESCs generating rosettes and lumens 70hrs after seeding. n = 3 biological replicates. Two-sided t test, **p<0.005 (K) Fluorescence arbitrary units of immunostained naïve markers NANOG and SOX2 in 3D in vitro lumens 70hrs after seeding. n = >300. Two-sided t-test; ***P = <0.0001

To examine how LIN28A-targeted mRNA decay coordinates pluripotency progression with epiblast morphogenesis, we modelled rosette formation and lumenogenesis using ESCs seeded in 3D hydrogel of basement membrane extract (BME) (3, 4) (Fig. 4F, S11A-B). After 48 hours in primed conditions, approximately half of ESC aggregates formed rosettes displaying apical–basal polarity of the actin cytoskeleton. About 10% of aggregates displayed a lumen, as monitored by live staining for the anti-adhesive sialomucin podocalyxin (PODXL), which repulses the apical membranes and lines the lumen, while the rest showed the apical exposure of PODXL in pre-apical patches (PrAPs) with no lumen yet visible. In contrast, approximately 40% of LIN28A KO ESCs aggregates formed rosettes, with the rest still forming unpolarized aggregates without lumens (Fig. 4G-H). After 70 hours, 85% of WT aggregates displayed an expanded lumen. In contrast, most KO aggregates remained unpolarized or formed rosettes proceeding only to PrAP formation, displaying only modest pre-apical patches of PODXL (Fig. 4I-J, S11B). Retention at the rosette stage was observed also by the expression of NANOG, SOX2 and the rosette marker OTX2 (3)(Fig. 4K, S11C-D), showing a failure in progression towards primed pluripotency. To examine the importance of the LIN28A cytoplasmic translocation for this transition, we generated LIN28A-GFP-NLS mESCs (Fig. S6A-B), in which LIN28A is restricted to the nucleolus. These had a similar phenotype as LIN28A KO, with impaired lumen formation and longer retention at the rosette stage (Fig. 4G-J). In conclusion, our findings show that in absence of the nuclear-to-cytoplasmic translocation of LIN28A, the morphogenetic events of the rosette are compromised and pluripotency transition arrests at the rosette stage. Nevertheless, epiblast epithelialization can alternatively be achieved even without rosette formation via an apoptosis-mediated pathway to clear non-polarized cells that lack basal lamina contacts (21). At that point multiple microlumens can form and apoptosis becomes necessary to merge them into a single cavity when all cells finally dismantled the naïve pluripotency network (Fig. 5A, S12A-B). The cells that retain naïve pluripotency do not require survival signals from the basal lamina and do not undergo apoptosis (21). We observed that LIN28A KO or LIN28A-NLS aggregates eventually formed multiple nascent lumens but, unlike WT cells, did not proceed with the cavity formation (Fig. 5A-B). For comparison, we used inducible Bcl-2 overexpression to inhibit apoptosis, which generated a similar phenotype without any effect on the differentiation propensity of ESCs, indicating that lumen multiplication is an extension of both delayed differentiation and lack of apoptotic clearance (Fig. 5A, S11A). In conclusion we demonstrated that the lack of the nuclear-to-cytoplasmic translocation of LIN28A prevents epiblast epithelialization via both the canonical pathway of membrane repulsion-driven lumenogenesis by the rosette, and the alternative pathway of embryonic cavitation.

**Figure 5.**
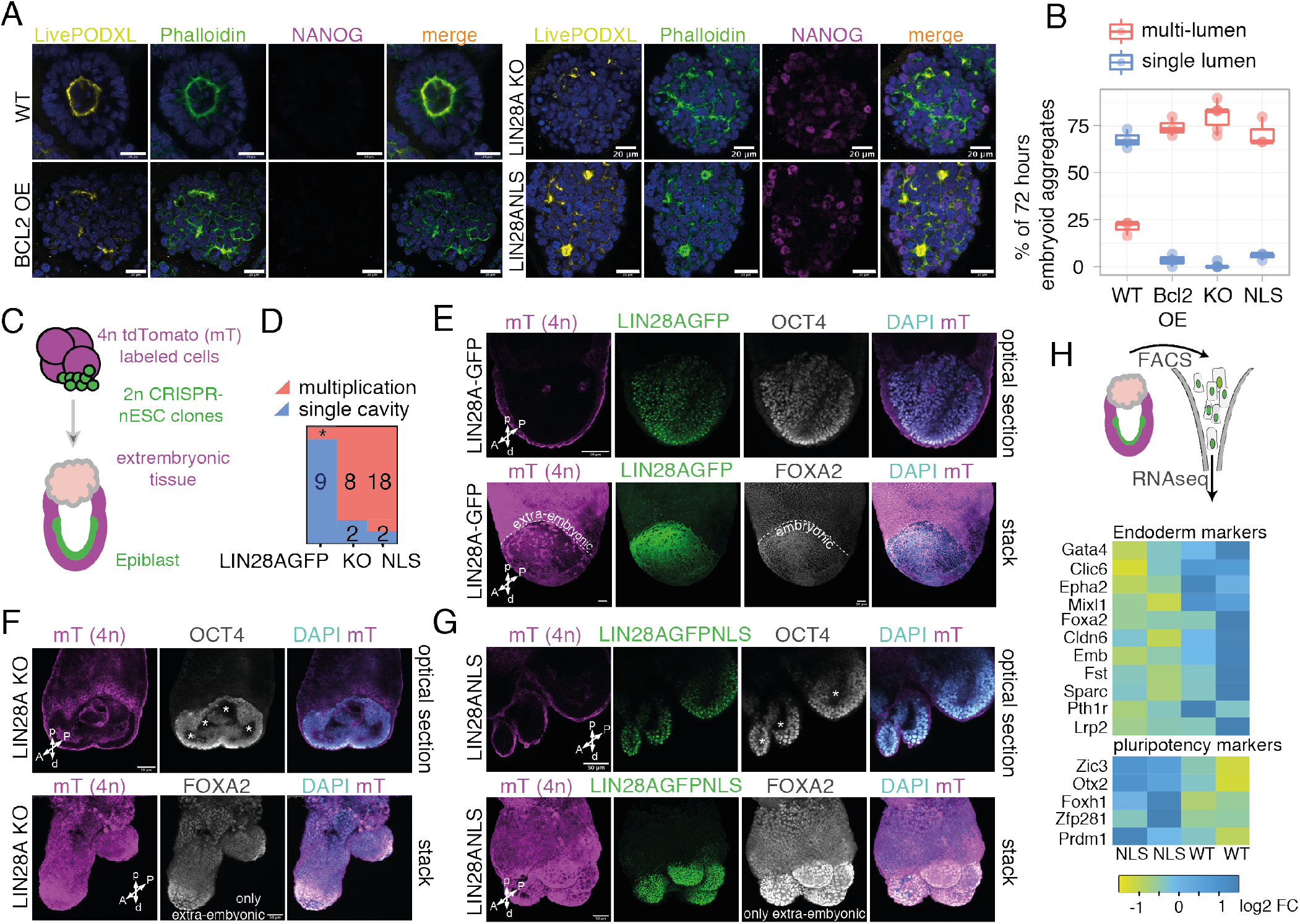
Loss of LIN28A or its nuclear retention causes an embryonic multiplication, multi-lumen formation and impedes gastrulation. (A) in vitro lumens formed after seeding WT, LIN28A KO and LIN28A-GFP-NLS ESCs in BME for 48hrs in 2iLIF followed by 70hrs of primed conditions, stained for indicated markers and (B) the percentage of ESCs generating single-or multi-lumens. (C) In vivo analysis of the developmental potency of ESCs exhibiting nuclear retention (LIN28AGFPNLS) or loss (KO) of LIN28A, compared to WT (LIN28AGFP) ESCs using tetraploid complementation assay, giving rise, respectively and exclusively, to embryonic and extraembryonic tissues. 4n extra-embryonic cells express membrane tdTomato (mT). (D) Distribution of embryo morphology and number of analysed embryos. *2 fused embryos with shared extraembryonic ectoderm (E-G) Shown are mESC-derived embryos (E7.5-E7.75) using (E) LIN28AGFP, (F) LIN28A KO and (G) LIN28A-GFP-NLS ESCs and representative analysis of mesendo-dermal (FOXA2) and epiblast (OCT4) marker by immunostaining (n = 10 for LIN28AGFP, n=20 using two independent clones of LIN28AGFPNLS, n = 10 for LIN28A KO). Blue, DAPI (nuclear stain). Scale bars, 50 um. A=anterior, P=posterior, p=proximal, d=distal. Dashed line indicates separation of visceral endoderm (extra-embryonic) and definitive endoderm (embryonic) that are both stained with FOXA2. * indicates multiple lumens (H) Heatmap showing up- and downregulated representative endoderm and pluripotency markers and genes determining left-right symmetry breaking, comparing the impact of LIN28AGFPNLS KO in FACS-sorted embryonic tissues of E7.5 tetraploid complemented embryos.

Lastly, we further assessed whether LIN28A-deficient embryos might still be able to achieve proper epithelialization of the epiblast at a later stage of embryogenesis. We generated completely mESC-derived embryos by tetraploid aggregation (16, 22). We aggregated the 4n 2-to 4-cell-stage embryos with 2n LIN28A-GFP, LIN28A-GFP-NLS or LIN28A KO ESCs (Fig. 5C-D, S13, S14A). Chimeric embryos derived from LIN28A-GFP mESCs generated morphologically normal egg cylinders (Fig. 5E). In contrast, the E7.5 to E8.0 embryos derived from LIN28A KO and LIN28A-GFP-NLS ESCs failed to form a central pro-amniotic cavity but generated multiple epiblast clusters, each containing a proto-pro-amniotic cavity (Fig. 5F, S14B). This validates our results from ESC-based embryogenesis systems (Fig. 4F-K, 5A-B) showing that, in the absence of cytoplasmic LIN28A translo-cation, local reorganization of the epiblast generates multiple lumens. These multiplied epiblasts retained OCT4 and SOX2 expression and failed to undergo gastrulation. This is exemplified by the lack of FOXA2 positive cells staining definitive endoderm in the embryonic tissues, and by the lack of visceral endoderm dispersal and therefore definitive endoderm formation (Fig. 5F-G). RNA sequencing of FACS-sorted GFP+ embryonic cells confirmed diminished expression of endodermal genes in LIN28A-GFP-NLS chimeras, and increased expression of pluripotency markers (Fig. 5H). Combined, these insights from mouse embryos show how an intrinsic mechanism of RBP translocation coordinates cellfate specification with morphological determination.

This work deciphers the molecular mechanism governing the first morphological transformation of epiblast cells, important for determining the failure or success of a pregnancy (2, 23). Notably, we find that selective mRNA decay dismantles the naïve pluripotency network in a manner that is coordinated with amniotic cavity formation. When LIN28A is ablated or trapped in the nucleus, embryos generate multiple incipient pro-amniotic cavities and epiblasts that retain naïve pluripotency features, which represents to our knowledge the first case of induced embryonic epiblast multiplication. Despite the well-studied role of LIN28A in let7 repression (18), our observed role of LIN28A in mRNA decay takes place before let7 induction, and is mediated by the direct binding of LIN28A to the 3’UTRs of destabilised mRNAs. A similar mechanism might also account for the let7-independent LIN28 roles in developmental timing of C.elegans larvae (24), mouse tissue repair (25), and human β cell differentiation (26). Given the central role of LIN28A-mediated mRNA stability in developmental progression, it likely contributes also to other functions of LIN28A in glucose metabolism (27) and tumorigenesis (19).

## ACKNOWLEDGEMENTS

The authors would like to thank George Daley and Kaloyan Tsanov for advice and generous reagent gifts. We thank Charlotte Capitanchik, Ira Iosuba for critical read-ing of the manuscript. For technical support we are grateful to Rahul Arora, Dorota Kurek, Timo Fisher, Matt Renshaw and other members of Crick science technology platforms for Advanced Sequencing, Light Microscopy and Flow Cytometry. This workwas supported bythe Wellcome Trust (Sir Henry Wellcome Fellowship to M.M., Senior Wellcome Trust Award to J.U. and N.M.L.), European Research Council (206726-CLIP and 617837-Translate to J.U.), Deutsche Forschungsgemeinschaft (DFG project 425470807 to H.Le.) and by the Francis Crick Institute, which receives its core funding from Cancer Research UK (FC001002), the UK Medical Research Council (FC001002), and the Wellcome Trust (FC001002). This preprint is formatted using a LATEX class template by Ricardo Henriques (CC BY 4.0) with minor alterations.

## AUTHOR CONTRIBUTIONS

M.M. conceived the study and performed the majority of the experiments and analysis apart from: 3D in vitro embryogenesis experiments were carried out and analysed by E.v.G, M.M., and supervised by D.t.B.. S.Sc. performed chimera experiments under the supervision of H.Li. M.M, C.M, T.K. and S.B. generated CRISPR lines, while M.M and J.R. performed imaging experiments and analysis with input from H.Le. M.M. and V.B performed and analysed mRNA-RBP occupancy experiments, while S.H. and J.M.P oversaw and performed MS workflow. I.rdl.M, M.M. and R.F. analysed the sequencing data and S.St. generated the machine learning workflow, with input from N.M.L, using CLIP datasets generated by M.M., F.L. and R. F. M.M. and J.U. jointly supervised the study. M.M., D.t.B. and J.U. wrote the manuscript.

---

**Figure S1:**
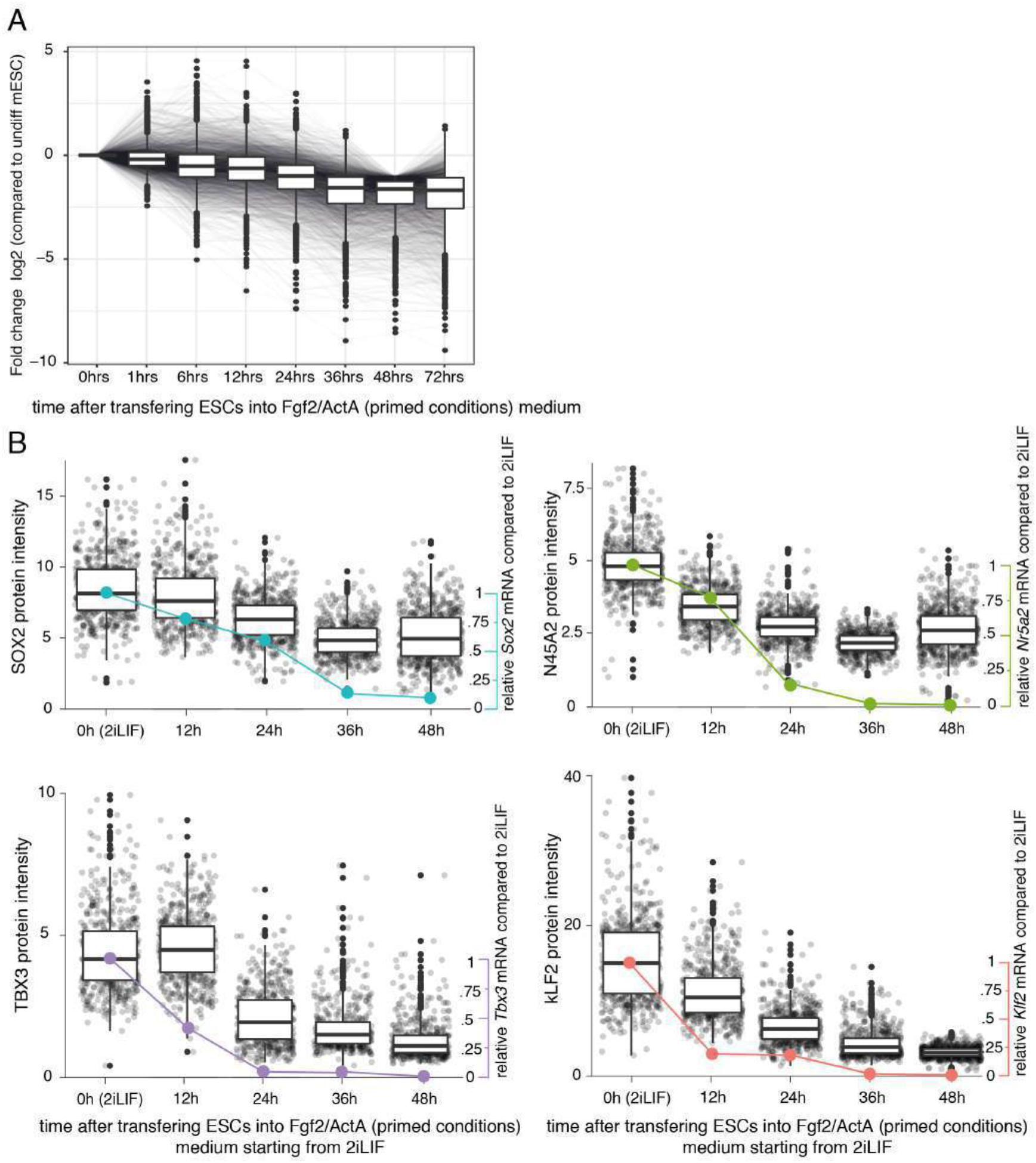
Earliest developmental gene and protein expression changes during naïve-to-primed transition. (A) Temporal dynamics of relative mRNA levels (compared to 2iLIF at Ohrs) of all genes downregulated during hours of ESC naive-to-primed differentiation. Error bars represent SEM. (B) Relative mRNA levels (compared to 2iLIF at Ohrs) and fluorescence arbitrary units of immunostained naive markers SOX2, KLF2, TBX3, NR5A2 (n = >200) during hours of ESC naive-to-primed differentiation. RNAseq data taken from (7).

**Figure S2:**
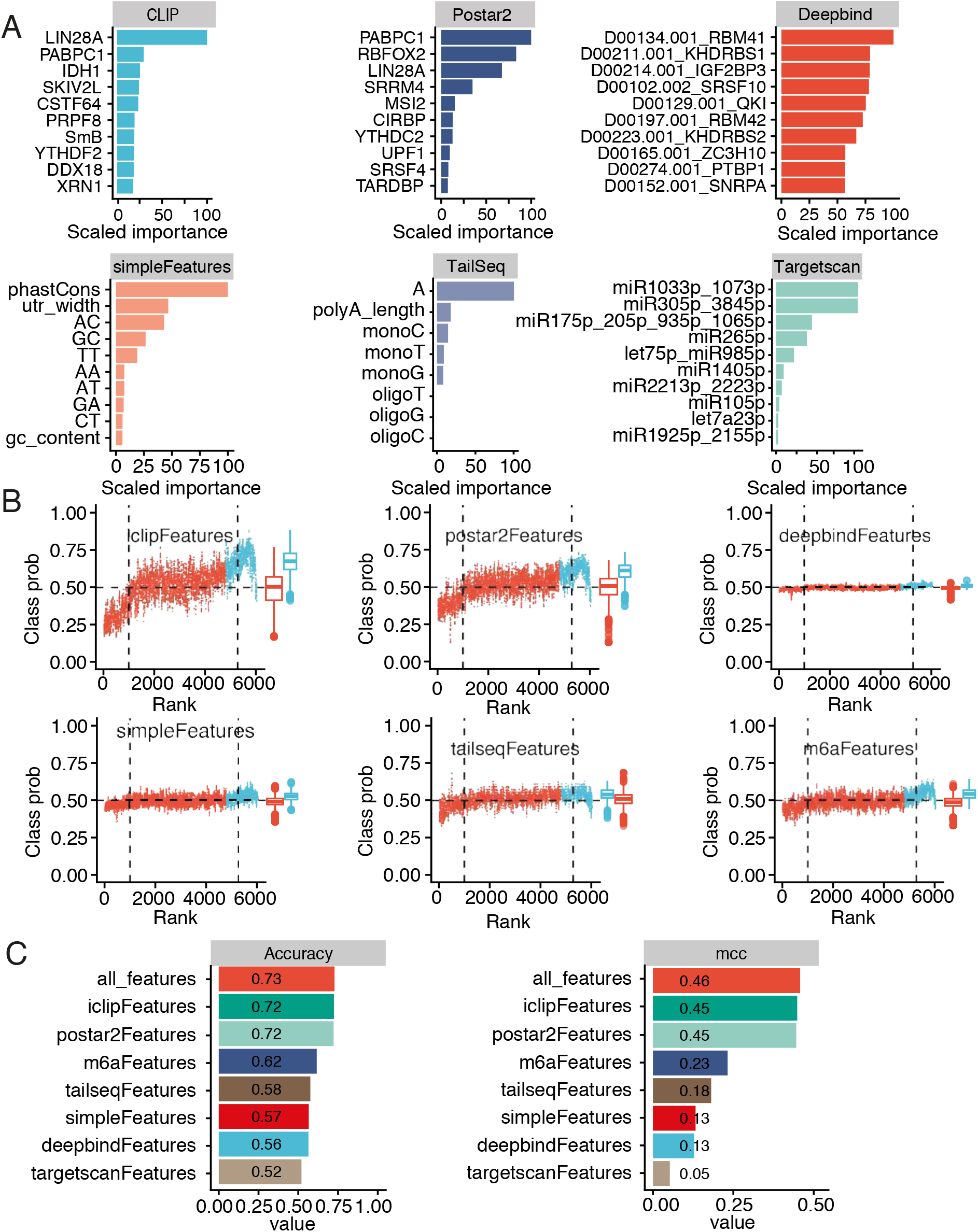
Feature importance impacting mRNA stability during naive-primed transition. (A) Feature importance ranking computed as relative influences, based on the relative influence, for the two best performing classifiers based on mESC CLIP, Postar2 CLIP-seq database, Deepbind, simple sequence features, TailSeq and TargetScan miRNA targets sites. (B) Ranking of all half-life chaning transcripts from UP (half-life higher in naive) towards DOWN (half-life higher in priming ESCs) compared with the class probability predicted with the binary model trained using features as presented in (A). Class probabilities were smoothed using the running mean (k=10). (C) Accuracy in predicting the change in half life of mRNAs during naive-to-primed conversion previously unseen by the model computed with features determined in (A). MCC = Matthews correlation coefficient.

**Figure S3:**
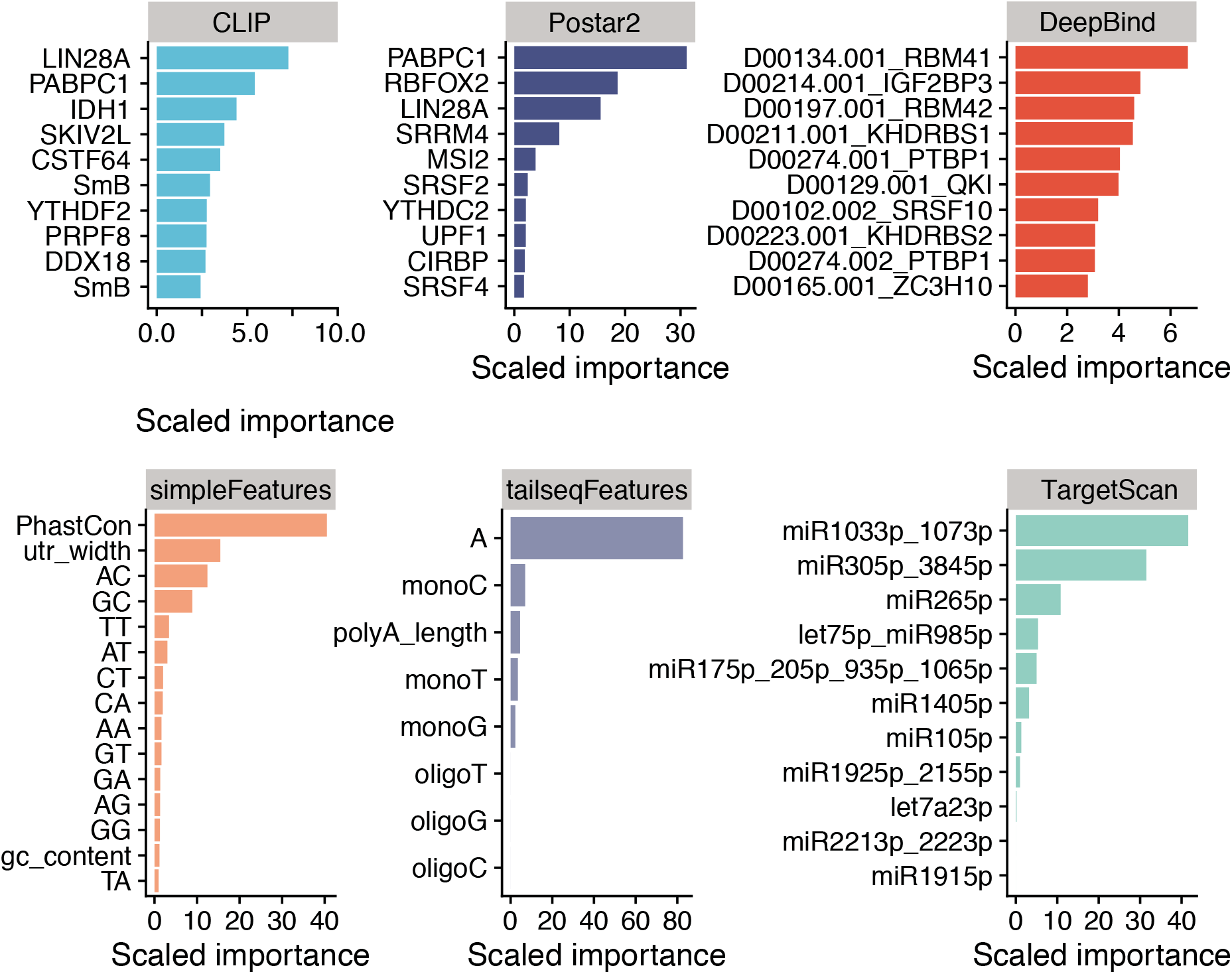
Permutation analysis of feature importance impacting mRNA stability during naive-primed transition. (A) Feature importance ranking computed as reduction in model performance by shuffling a particular feature. based on the relative influence, for the two best performing classifiers based on mESC CLIP, Postar2 CLIP-seq database, Deepbind, simple sequence features, TailSeq and TargetScan miRNA targets sites.

**Figure S4:**
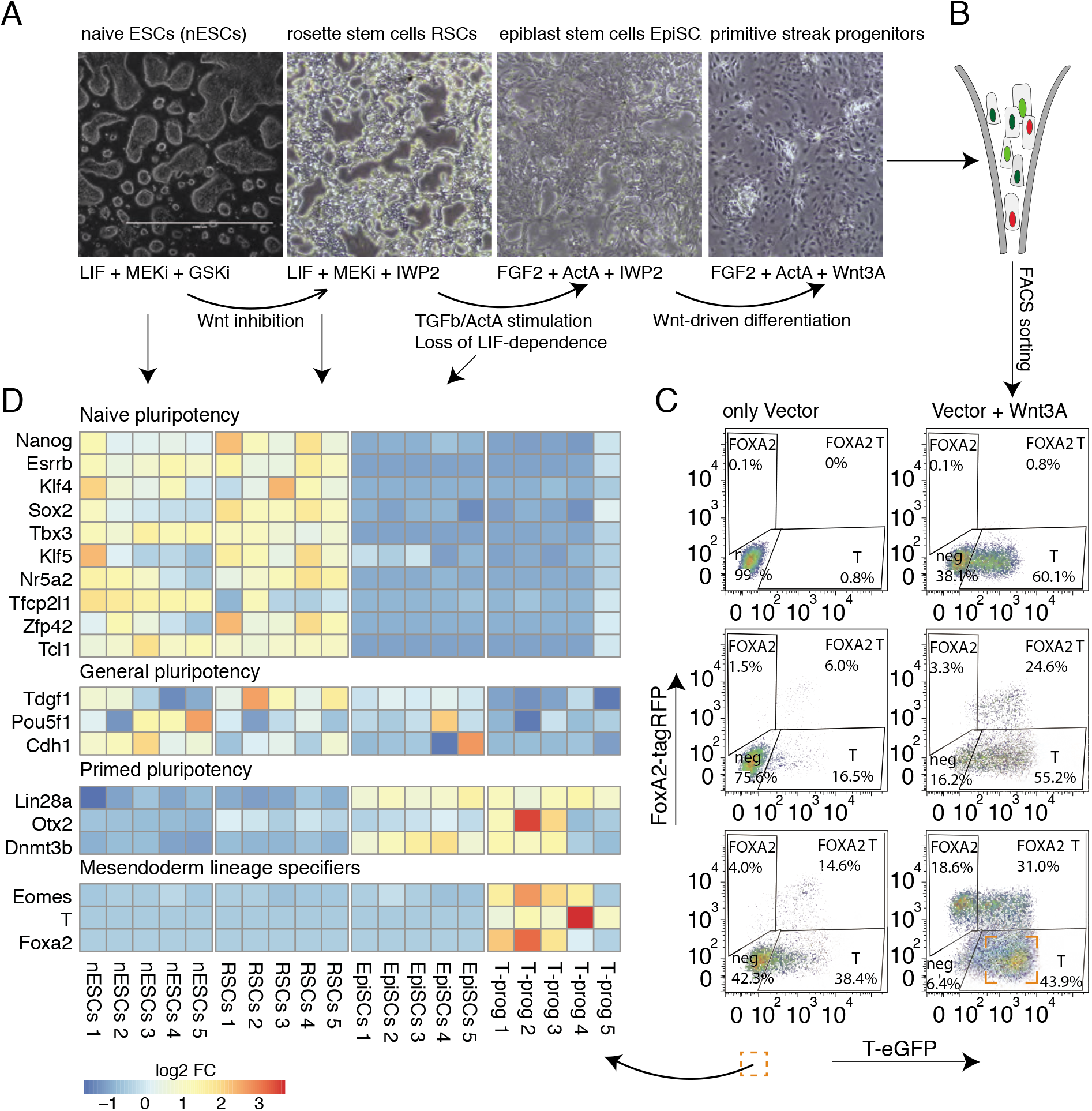
Benchmarking and isolation of homogeneous population of nESCs, rosette stem cells, epiblast stem cells (EpiSCs) and FACS-sorted mesodermal progenitors. (A) Representative photomicrographs of naive mESCs, rosette stem cells, EpiSCs and 3days-differentiated EpiSCs with indicated morphogen changes, which instruct transitions of ESCs from naive, rosette, epiblast towards mesoderm progenitors. Primed Wnt-inhibited EpiSCs respond directly to germ layer induction, allowing us to enrich for the earliest differentiated mesoderm progenitors by concomitantly FACS-sorting the mesoderm progenitors expressing T (Brachury) reporter in T-GFP FoxA2-tagRFP ESCs upon 72hrs Wnt3a treatment. (B) Flow cytometry schematic and (C) diagrams of dual reporter Foxa2-RFP T-eGFP EpiSCs treated with Wnt3a and analyzed according to the relevant fluorescent emissions. Cells were maintained in the presence of IWP2 prior to the experiment and vector stands for bFGF- and ActA-containing basal medium. (D) Expression heat map of representative naïve pluripotency factors (Nanog, Esrrb, Klf4, Sox2, Tbx3, Klf5, Nr5a2, Tfcp2l1, Zfp42, Tcl1), core general pluripotency network (Tdgf1, Pou5f1/Oct4, Cdh1), early priming factors (Lin28a, Otx2, Dnmt3b) and lineage specifiers (Eomes, T, Foxa2).

**Figure S5:**
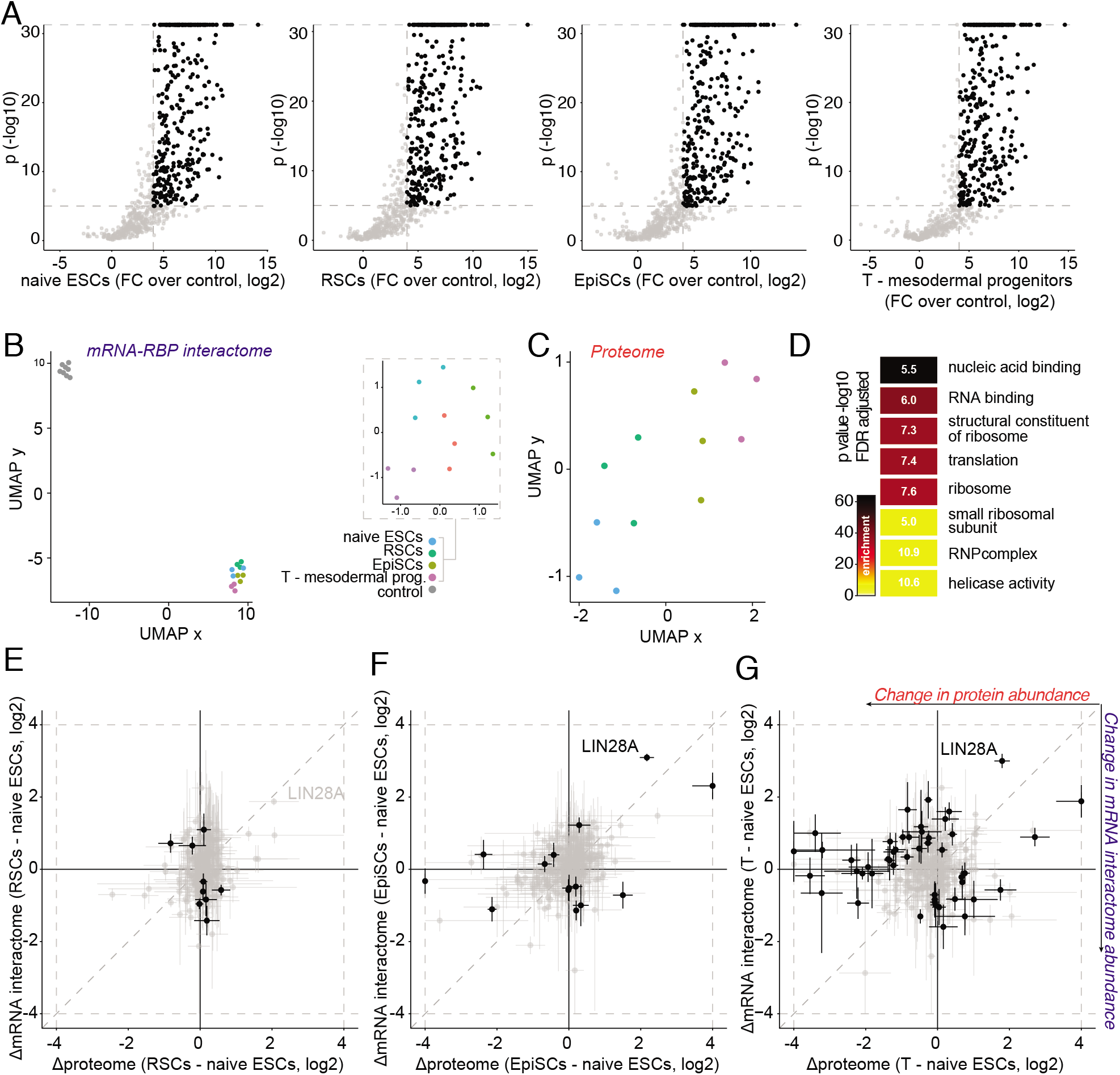
Developmental mRNA-RBP interactome capture identifies RBPs that dramatically change mRNA binding during early development. (A) Volcano plots, depicting lasso based p-value (-log10) and log2 fold-change, from enrichment analysis of mRNA-interactome capture in indicated conditions over controls. Proteins, considered significantly enriched in individual conditions, are highlighted in black. Significance cut-offs are indicated by dotted line. Proteins, enriched in all conditions, were designated bona-fide RBPs and used for analysis of dynamic binding across conditions. (B) UMAP dimensionality reduction applied to mRNA-interactome samples and controls, and separately to non-control samples as indicated, and (C) to proteome samples. (D) Top 8 significant GO-terms, enriched in a set of candidate RNA-binding proteins over all proteins, detected in the MS analysis. (E, F, G) Scatter plots depicting gaussian protein abundance changes (+/− gaussian standard deviation) in mRNA-interactome versus proteome between rosette stem cells and nESCs (E), EpiSCs and nESCs (F) and mesodermal progenitors and nESCs (G). Discrepancy between changes was analysed using lasso-based linear model and significant proteins with absolute log2 discrepancy > 0.25 and p < 0.001 were considered significant and are highlighted in black. The values are truncated for visualisation purposes according to the dotted frame.

**Figure S6:**
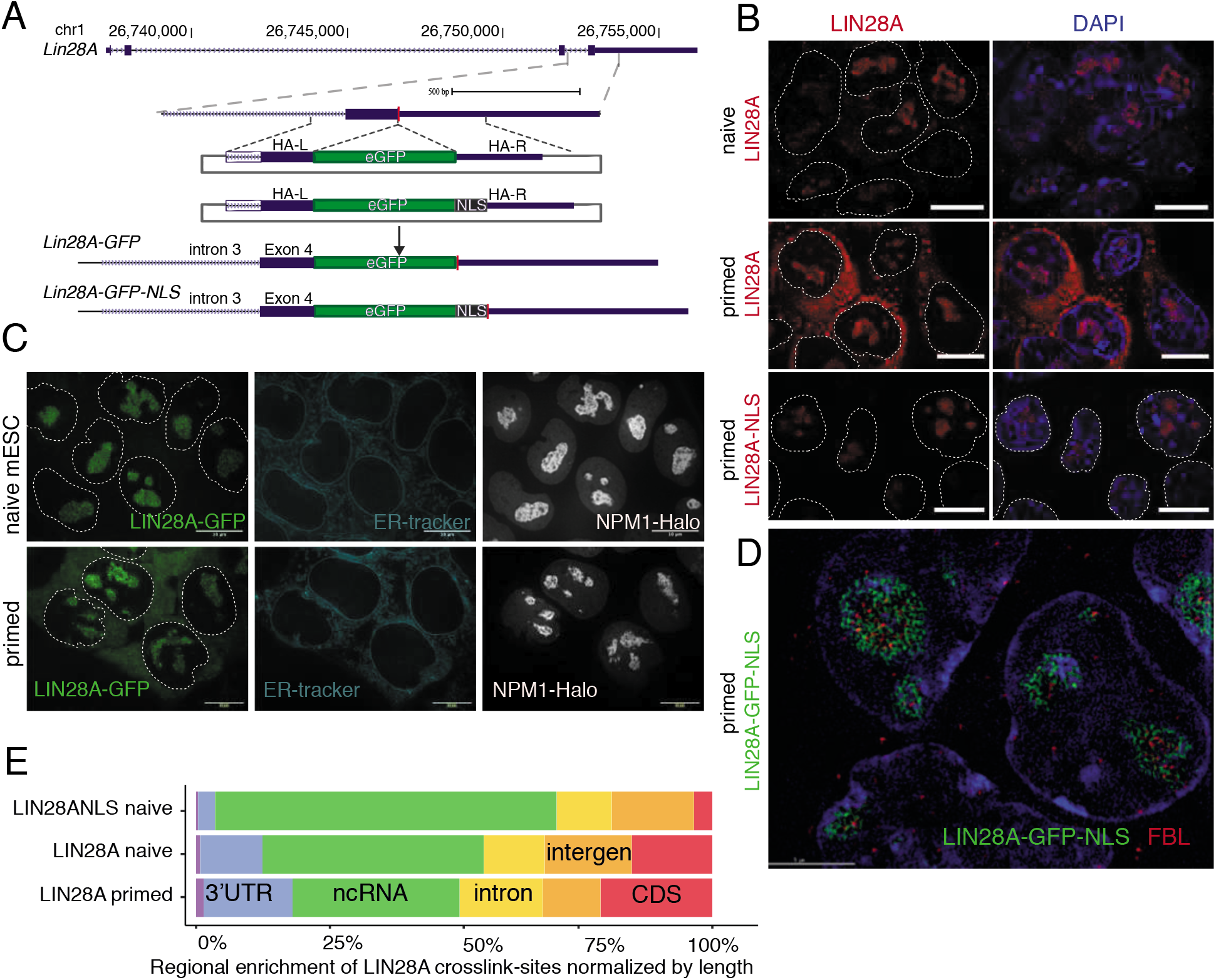
Confirmation of LIN28A localisation pattern using live imaging and immunofluorescence. (A) Schematic overview of eGFP or eGFP-NLS insertion into the Lin28a locus via CRISPR/Cas assisted gene editing. Target site followed by PAM motif (5’-AGG) is located on the top strand (positive strand) upstream of LIN28A stop codon. Integration is facilitated by double strand breaks created by Cas9 directed to the target sequence by a specific gRNA. (B) Live cell representative fluorescence micrographs of naive ESCs and primed ESCs expressing the endogenous LIN28A-GFP in live cells in 2iLIF and upon 16hrs of transfering ESCs to the primed conditions, coinciding with the time point of Fig. 1B. Scale bar: 10μm. (C) Representative immunofluorescence photomicrographs or naive ESC and primed EpiSC WT cells and primed LIN28A-NLS ESCs. Blue, DAPI (nuclear stain). Scale bars, 10 μm. (D) Fluorescent structured illumination micrograph (SIM) of primed LIN28A-GFP-NLS cells showing exclusive nuclear localisation of LIN28A-NLS enriched in the nucleolar compartment as determined by FBL staining. Blue, DAPI (nuclear stain). Scale bars, 5 μm. (E) Distribution of binding sites per RNA type and mRNA region as defined by mapping of iCLIP unique cDNAs. Presented are average values of 3 iCLIP replicates per naive LIN28AGFP ESCs, primed LIN28AGFP ESCs and naive LIN28AGFP-NLS ESCs.

**Figure S7:**
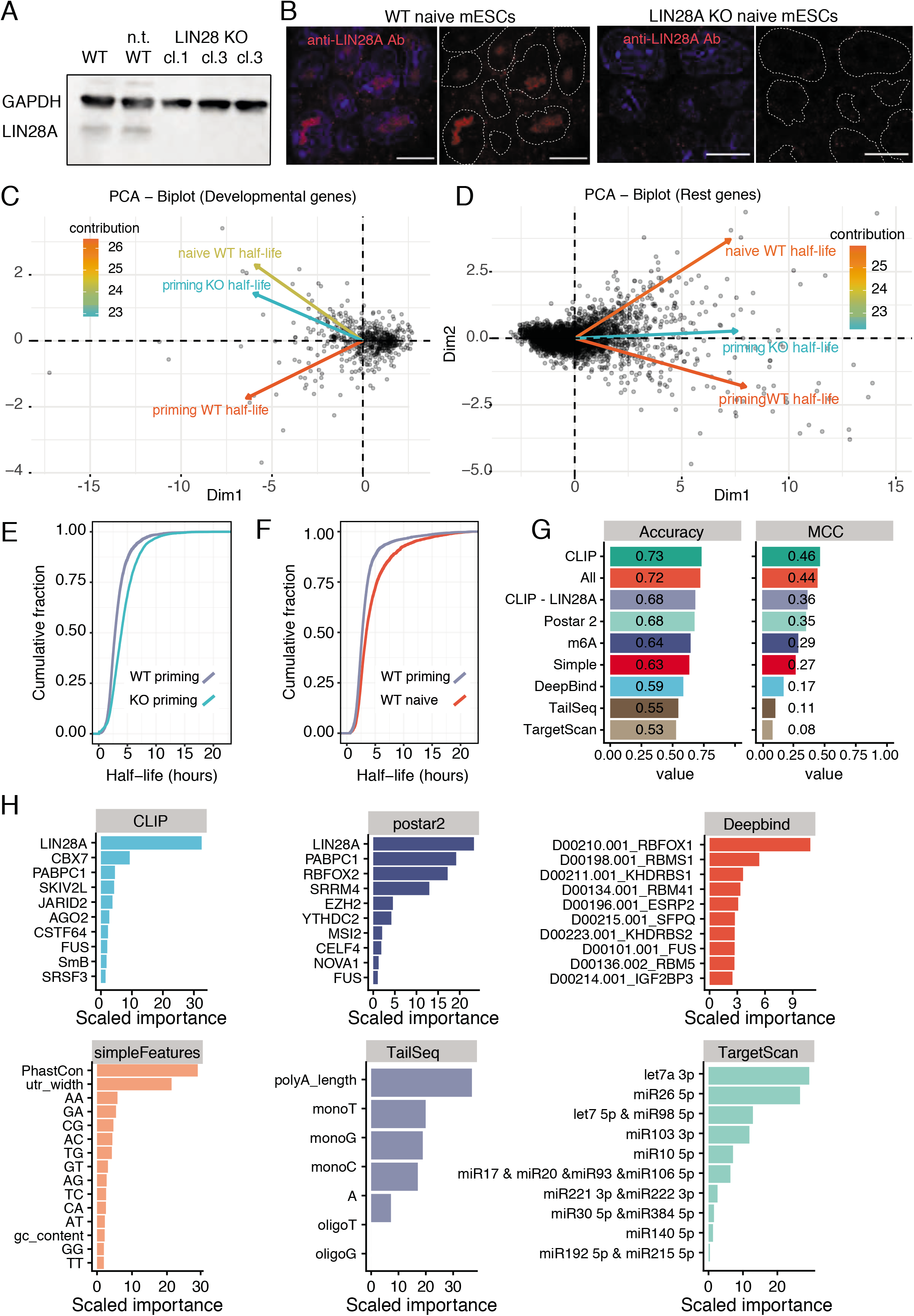
Feature importance and analysis of genes impacting mRNA stability upon LIN28A KO. (A) Western blot analysis of LIN28A and histone H3 in LIN28A WT mESCs, non-targeted LIN28A clone and 3 independent verified LIN28A KO clones. (B) Representative iSIM immunofluorescence photomicrographs or naive WT and LIN28A KO ESCs, validating the loss of LIN28A protein. (C-D) BiPlot of principal component analysis of mRNA half-live identified by SLAM-seq during naïve-to-primed pluripotency transition (naïve ESCs towards priming ESCs). Each black dot represents the covariance decomposition of each gene in the first and second dimension. On the left developmental genes (C) and the rest of the genes on the right (D). Arrows display the eigenvector direction for each experiment and the colour indicates their contribution to the model. (E-F) Cumulative distribution of mRNA stability changes in primed LIN28A KO relative to primed wild-type ESCs (E) and naive relative primed WT ESCs (F). (G) Accuracy in predicting the change in half life of mRNAs upon LIN28A KO previously unseen by the model computed with features determined in (H). MCC = Matthews correlation coefficient. (H) Feature importance ranking predicting mRNA half-life changes between LIN28A WT and KO priming ESCs based on the relative influence, for the two best performing classifiers based on mESC CLIP, Postar2 CLIP-seq database, Deepbind, simple sequence features, TailSeq and TargetScan miRNA targets sites.

**Figure S8:**
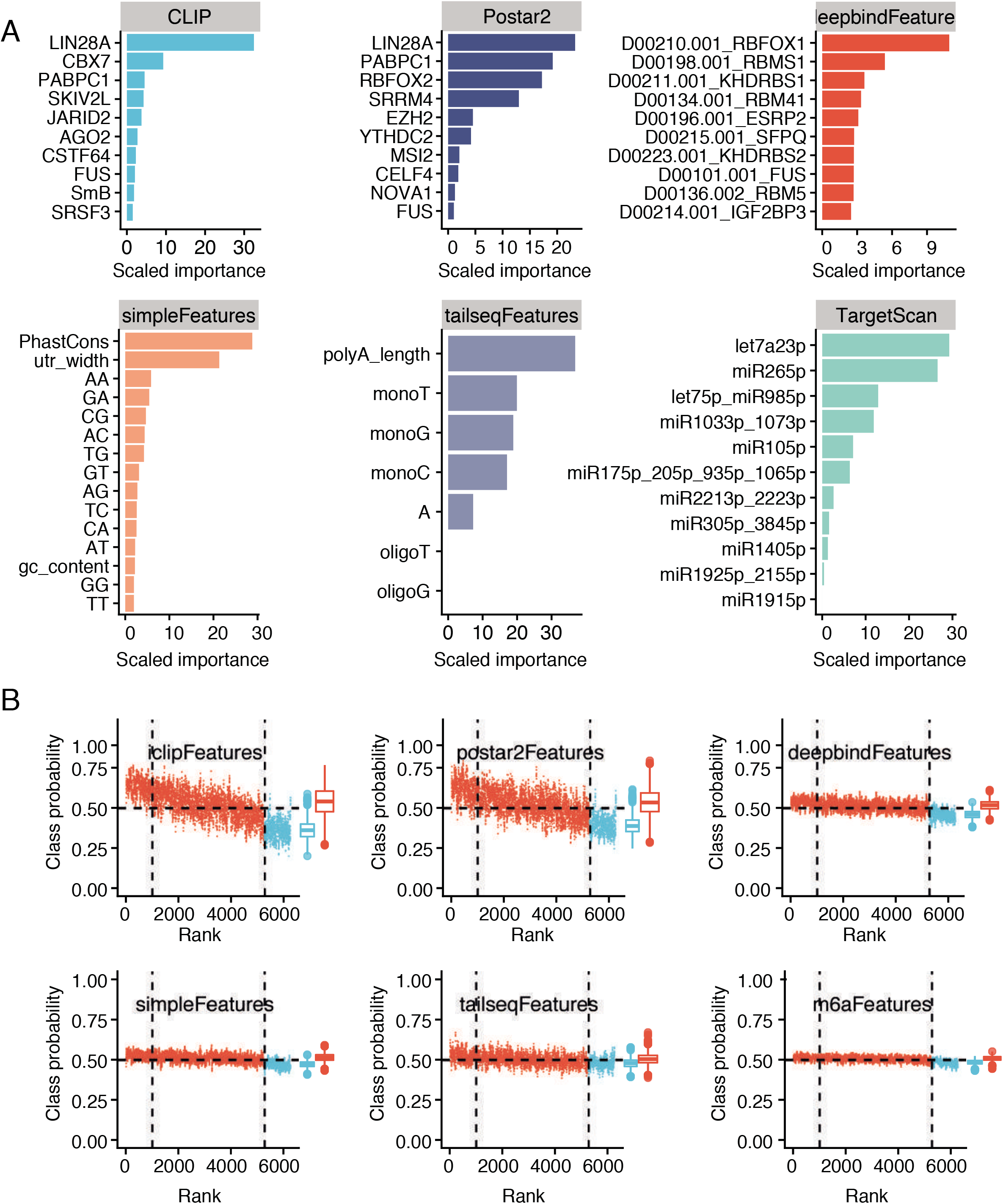
Permutation analysis of feature importance impacting mRNA stability upon LIN28A KO. (A) Feature importance ranking computed as reduction in model performance by shuffling a particular feature. based on the relative influence, for the two best performing classifiers based on mESC CLIP, Postar2 CLIP-seq database, Deepbind, simple sequence features, TailSeq and TargetScan miRNA targets sites. (B) Ranking of all half-life changing transcripts from UP towards DOWN upon LIN28A KO compared with the class probability predicted with the binary model trained using CLIP, Postar2, deepbind, simple, tailseq and m6A features. 1000 most destabilized transcripts are labeled blue. Class probabilities were smoothed using the running mean (k=10).

**Figure S9:**
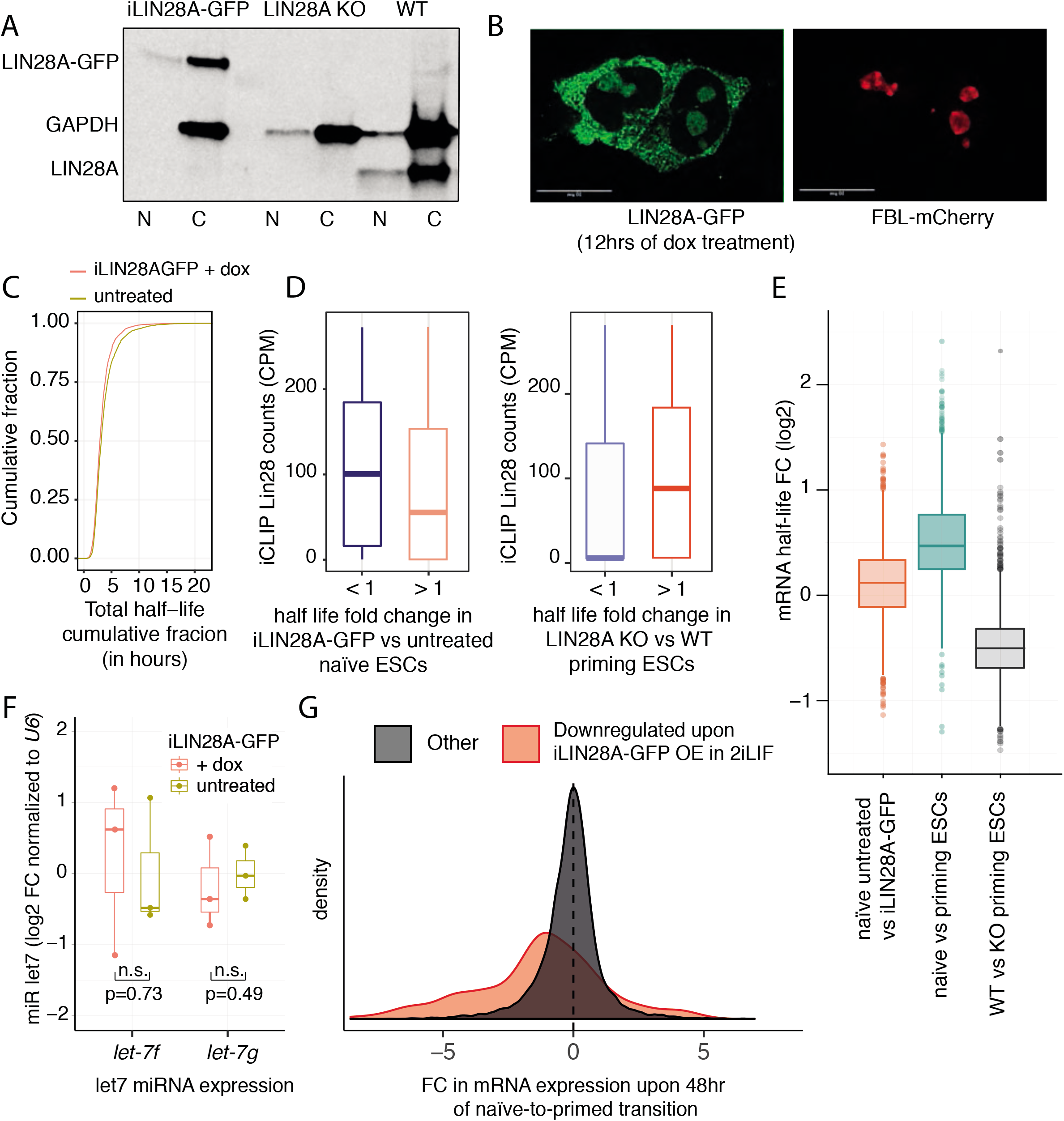
mRNA stability and expression changes upon inducible LIN28A-GFP overexpression in 2iLIF. (A) Western blot analysis of LIN28A, LIN28A-GFP and GAPDH in wild-type primed EpiSCs, doxycyclin-treated and -untreated iLIN28A-GFP naïve ESCs. (B) Live cell representative fluorescence micrographs of naive ESCs expressing endogenous FBL-mCherry and upon dox-inducible LIN28A-GFP overexpression in 2iLIF, validating the cytoplasmic accumulation of LIN28A-GFP protein. (C) Cumulative distribution of mRNA stability changes in naïve dox-treated iLIN28A-GFP relative to its untreated control. (D) Normalized LIN28A binding (y-axis) to 3’UTRs of transcripts according to change in their mRNA half-life upon iLIN28A-GFP overexpression in naive ESCs (left) and upon LIN28A KO in priming ESCs (right). (E) Box-plots outlining the extent and direction of global mRNA half-life fold change upon iLIN28A-GFP induction in naive ESCs, upon embryonic priming and upon LIN28A KO in primed ESCs. (F) Boxplot depicting relative levels of miR let-7 in naïve iLIN28A-GFP mESCs with and without doxycyclin treatment (n = 3 independent replicates, two sided paired t test values indicated). (G) Density of fold-change in gene expression changes of transcripts downregulated upon 48hrs of transferring ESC cells into primed medium(7). Genes that are downregulated upon 6hrs of dox-induced iLIN28A-GFP overexpression are labled in red, while remaining genes are labeled black.

**Figure S10:**
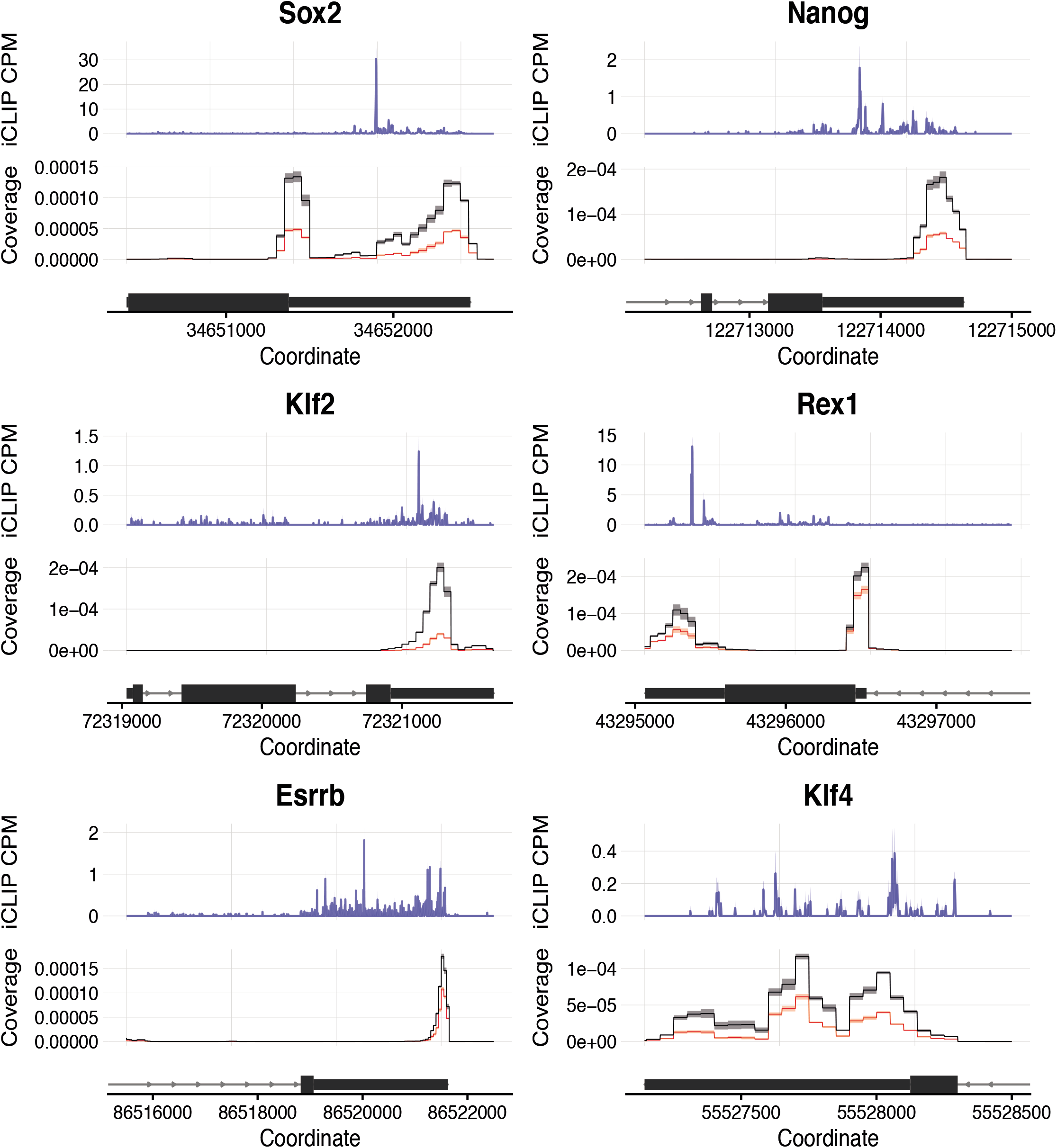
Representative LIN28A crosslinking to naive pluripotency transcripts and changes in their mRNA expression. Representative LIN28A-GFP crosslinking positions in the 3’UTRs of naive pluripotency transcripts (blue) and the frequencies of QuantSeq mapped to the transcripts encoding naive pluripotency factors comparing untreated (black) and dox-induced (6hours, red) iLIN28A-GFP naive ESCs.

**Figure S11:**
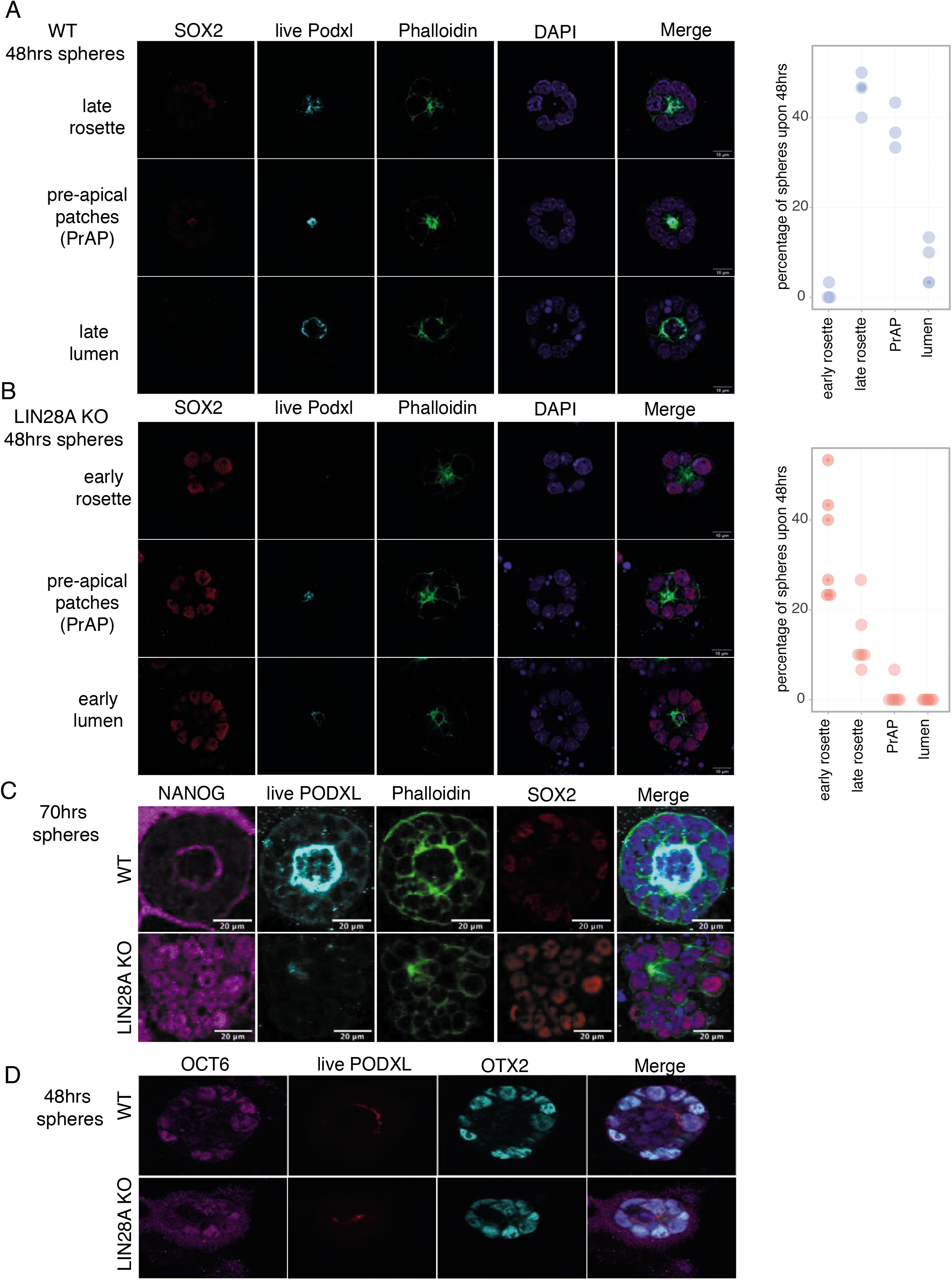
Representative examples and quantifications of rosette, pre-apical patches and late lumens. (A) Rosettes and in vitro lumens 48hrs after seeding wild-type ESCs in BME in priming conditions, stained for indicated markers and the percentage of ESCs generating rosettes, pre-apical patches and lumens 48hrs after seeding. n = 3 biological replicates. (B) Rosettes and in vitro lumens 48hrs after seeding LIN28A KO ESCs in BME in primed conditions, stained for indicated markers and the percentage of ESCs generating rosettes, pre-apical patches and lumens 48hrs after seeding. n = 3 biological replicates. (C) Lumens 70hrs after seeding wild-type and LIN28A KO ESCs in BME in primed conditions, stained for indicated markers. (D) Aggregates 48 h after seeding ESCs in BME in primed conditions, stained for rosette marker (OTX2) and marker of primed pluripotency (OCT6).

**Figure S12:**
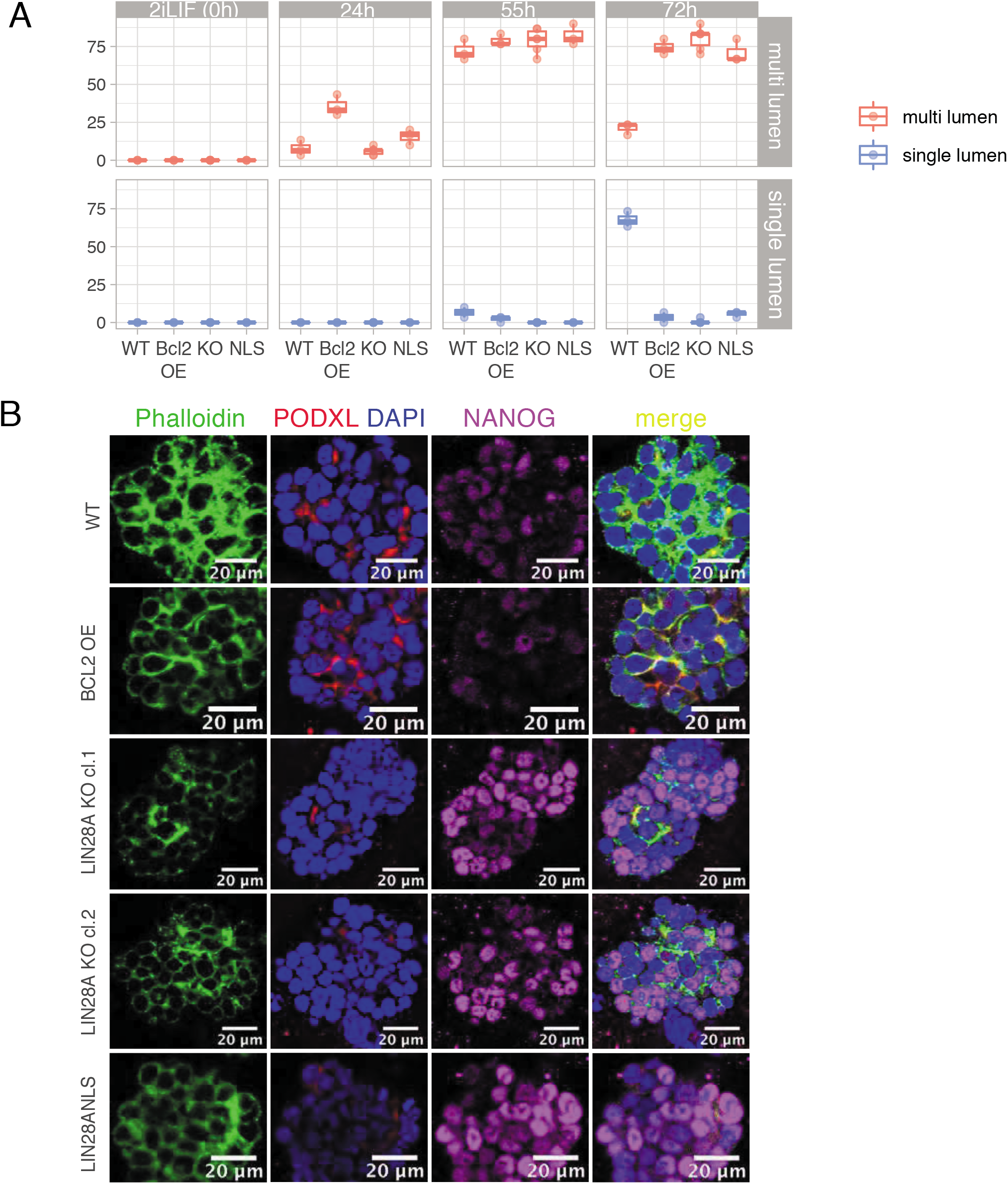
Representative examples and quantifications of cavitation-induced multi-lumen formation. (A-B) 3D large aggregates formed after seeding WT, LIN28A KO and LIN28A-GFP-NLS ESCs in BME for 48hrs in 2iLIF followed by the indicated length of culturing in primed conditions. (A) Aggregates were stained for indicated markers upon transferring to primed culturing conditions for 72hrs (as presented in Fig. 5A) and upon 55hrs (B). This experiment assesses whether cell death of primed cells is required for a unified single lumen formation in large self-organising aggregates. Suppression of apoptosis by overexpressing the pro-survival gene Bcl-2 impairs embryonic cavitation and gives rise to multiple incipient lumens without impacting differentiation propensity of ESC, while in contrast, the effect of loss of cytoplasmic LIN28A is a result of defective differentiation, as seen by retained expression of naive pluripotency markers, whereas Bcl-2 overexpression had no effect on the differentiation propensity of ESCs.

**Figure S13:**
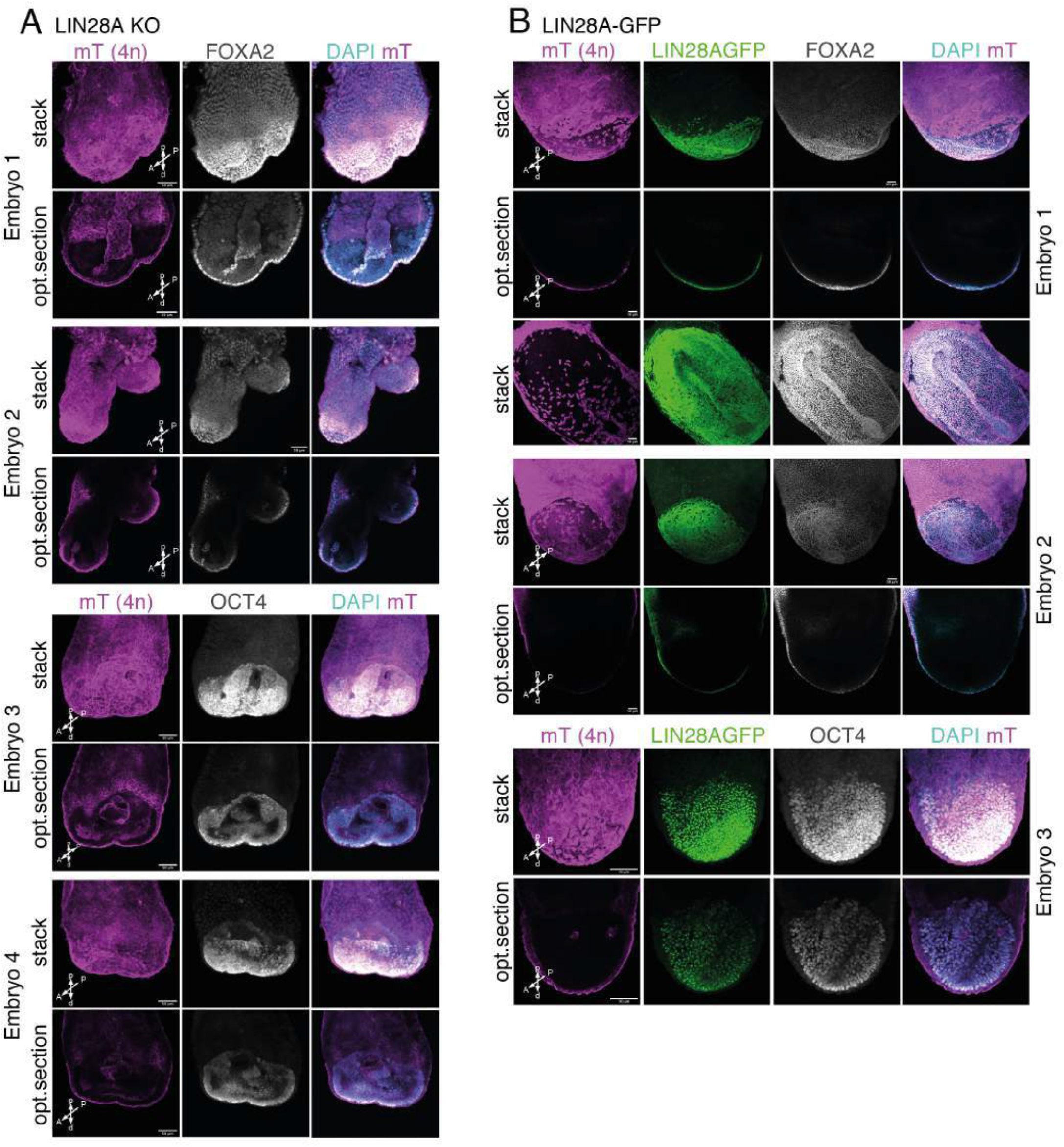
The developmental competence and multi-lumen formation of LIN28A KO embryos compared to control LIN28A-GFP embryos. (A-B) Representative examples of mouse embryos (E7.5-E7.75) resulting from aggregations of (A) LIN28AKO, and (G) LIN28A-GFP ESCs with 4n 2-to-4-cell-stage embryos and representative analysis of endodermal (FO×A2) and epiblast (OCT4) marker by immunostaining (n = 10 for LIN28AGFP, n=2O using two independent, n = 10 for LIN28A KO). Blue, DAPI (nuclear stain). Scale bars, 50 urn. A, anterior; P, posterior; p, proximal; d, distal.

**Figure S14:**
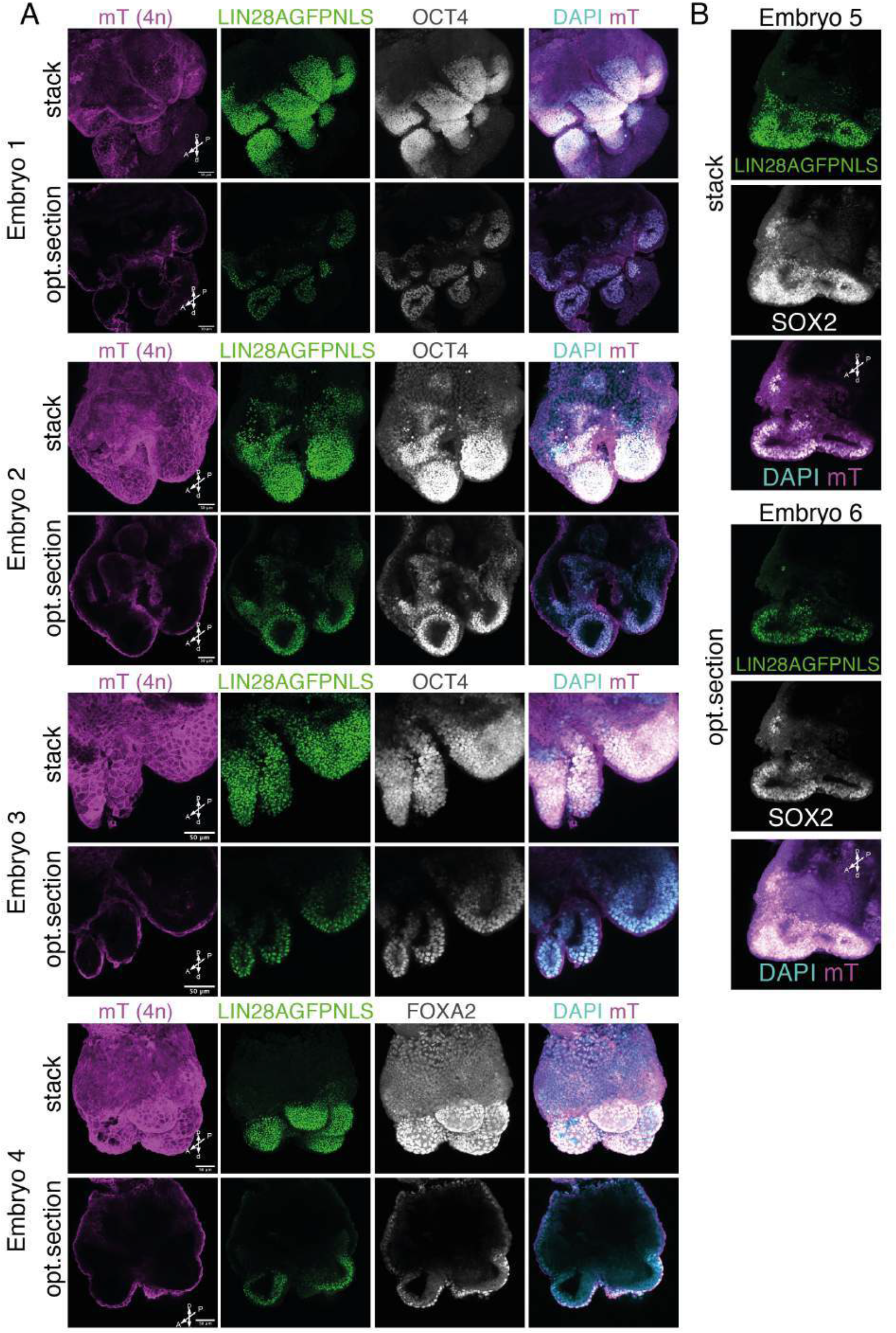
The developmental competence and multi-lumen formation of LIN28A-GFP-NLS embryos. (A-B) Representative examples of mouse embryos (E7.5-E7.75) resulting from aggregations of LIN28A-GFP-NLS ESCs with 4n 2-to-4-cell-stage embryos and representative analysis of (A) endodermal (FOXA2) and epiblast (OCT4) marker and (B) naive pluripotency marker (SOX2) by immunostaining (n=2O using two independent LIN28A-GFP-NLS clones). Blue, DAPI (nuclear stain). Scale bars, 50 urn. A, anterior; P, posterior; p, proximal; d, distal.

## Material and Methods

### Embryonic stem cell culture

All mouse ESC lines e.g., J1 IDG3.2 and V6.5, were maintained in 1:1 Neurobasal (21103049) DMEM (11320033) medium containing N2 (17502048) and B27 (17504044) supplements, 1% Glutamax (35050), 1% nonessential amino acids (1140050), 0.1 mM 2-mercaptoethanol (31350-010) (all Thermo Fisher Scientific), 1 mM MEK inhibitor PD0325901 (1408, Axon Medchem), 3 mM GSK3 inhibitor CHIR99021 (4953/50 Tocris), and 1,000 U/ml LIF (ESGRO ESG1107, Merck), a condition named 2iLIF. Cells were passaged using Stempro-Accutase (A1110501, Thermo Fisher Scientific). Cells were grown on gelatin coated plates which were prepared by first incubating the wells 0.1% gelatin (EmbryoMax® ES-006-B, Merck) with 1% FBS (EmbryoMax® ES Cell Qualified FBS, ES-009-B, Merck) for 1 h at 37 °C followed by a rinse with PBS.

To transit from naive ESC to priming ESC cells, accutased naive ESCs were seeded onto gelatine and FBS-coated plates in N2B27 medium supplemented with recombinant 20 ng ml–1activin A (338-AC-050, R&D Systems) and 12 ng ml–1 bFGF (100-18B, Peprotech) for 12 hours or as indicated in the Figure legends (12, 24, 36 or 48 hours). To determine developmental genes that are downregulated during naive-to-primed transition, the publicly available data were used (Yang et al., 2019a) and analyzed as described in the RNAseq analysis section. Primed epiblast stem cells (EpiSCs) were cultured on gelatine and FCS-coated plates in N2B27 medium supplemented with recombinant 20 ng ml–1activin A (338-AC-050, R&D Systems) and 2 μM IWP2 (3533, Tocris) to suppress spontaneous differentiation. EpiSCs were passaged 1:4–1:10 every 3 days by triturating the colonies into small clumps using 0.5 mg ml–1 collagenase IV (Sigma). MEK/WNT-inhibited RSCs were cultivated with 1,000 U ml–1 LIF, 2 μM IWP2, 1 μM PD (manufacturers as mentioned above) and were passaged as ESCs.

To induce primitive streak differentiation we utilized a double reporter mESC line modified to express distinct fluorescent proteins regulated by the promoters of T (Brachyury) and Foxa2 genes, where the expression of T and Foxa2 alone is indicative of mesodermal and endodermal commitment, respectively (Burtscher and Lickert, 2009). First, EpiSCs from these cells were derived by transiting ESCs to primed cells with 1,000 U ml–1 LIF, 2 μM IWP2, 20 ng ml–1 Activin A and 12 ng ml–1 bFGF (manufacturers as mentioned above). After 3 days, LIF has been omitted and derived EpiSCs can be passaged 1:4–1:10 every by triturating the colonies into small clumps using 0.5 mg ml–1 collagenase IV (Sigma). To derive mesodermal progenitors utilizing the T-eGFP FoxA2-tagRFP double knock-in reporter cell lines, the IWP2 that is in EpiSCs used to suppress differentiation was replaced by 250ng/mL of WNT3A (in house produced as described previously (Neagu et al., 2020)) and fresh medium was applied daily. Expression of T-eGFP and FoxA2-tagRFP, indicative of mesodermal and endodermal progenitors, was monitored by Flow cytometry. All cell lines in culture were tested for mycoplasma contamination every 2–3 months.

### Stem Cell-based embryogenesis

ESCs were triturated to a single cell suspension using 0.05% trypsin–EDTA (25200056, Life Technologies) and washed with PBS. Cells were resuspended in ice-cold BME (BME 2 RGF, AMSBIO, 3533-010-02) at a density of 3,000 cells per μl. Drops of 20 μl were in a 8-chamber slide (80826, Ibidi) and incubated for 5 min at 37 °C until the BME solidified. The plate was then filled with prewarmed N2B27 medium supplemented with 1.000 U/ml LIF, 20 ng/ml activin A, 12 ng/ml bFGF (both Peprotech) and 2 μM IWP2 (681671, Merck) and cultured at 37 °C and 5% CO_2_ for indicated times. N2B27 consisted of one volume DMEM/F12 combined with one volume Neurobasal medium supplemented with 0.5% N2 supplement, 1% B27 supplement, 0.033% BSA 7.5% solution, 50 μM β-mercaptoethanol, 2 mM Glutamax, 100 U/ml penicillin and 100 μg/ml streptomycin (all from Invitrogen). For the increased sized spheres, 1,500 cells per μl were used and cells were first cultured in 2iLIF conditions for 48hours, followed by primed induction in N2B27 supplemented with LIF, bFGF, activinA and IWP2 for indicated times.

For live PODXL stainings, live cultures were incubated for 1h in a cell culture incubator with culture medium containing 1:500 anti-PODXL antibody (MAB1556, R&D Systems). The cells and BME drops were subsequently fixed with 4% PFA, pre-warmed to 37 °C, for 30 min. For immunostaining, the fixed cells were permeabilized with 0.3% Triton/0.1 M glycine in PBS for 30 min at room temperature, washed with PBS containing 0.05% Tween 20 (PBST) for 20 min, incubated in blocking buffer for 1 h (1% BSA, 5% donkey serum in PBST) and incubated overnight at room temperature in blocking buffer with the primary antibodies 1:1000 Sox2 (GT15098, Immune Systems) or 1:250 Nanog (rec-rcab0002p-f, Coso Bio). The next day, cells were washed three times for 30 min with PBST before incubation for 2 h with secondary antibodies in blocking buffer. After three washes with PBST for 30 min, phalloidin was stained with 1:200 phalloidin-594 (A12381, Invitrogen) in PBST for 1.5 h. Nuclei were stained with DAPI (Molecular Probes) for 30 min at room temperature and washed three times with PBS. The stained preparations were covered with Prolong Gold (P36930, Invitrogen) and imaged using a Leica SP8 inverted laser-scanning confocal microscope.

### Generation of CRISPR/Cas9 genome engineering mESC lines

For the generation of *Lin28a-GFP* or *Lin28a-GFP-NLS* J1 ESCs, specific gRNAs targeting upstream of the stop codon (Supplementary Table) were cloned into a modified version of the SpCas9-T2A-GFP/gRNA plasmid (px458172, Addgene plasmid #48138), where we fused a truncated form of human Geminin (hGem) to SpCas9 in order to increase homology-directed repair efficiency generating SpCas9-hGem-T2A-GFP/gRNA. To generate *Lin28a-GFP* or *Lin28a-GFP-NLS* targeting donors, dsDNA gene fragments were synthesized and cloned into a vector carrying ~400 bp homology arms (IDT, Coralville, IA, USA) (Supplementary Table). For targetings, wild-type ESC were transfected with a 4:1 ratio of donor oligo and SpCas9-hGem-T2A-GFP/gRNA construct. Positively transfected cells were isolated based on GFP expression using fluorescence-activated cell sorting (FACS) and plated at clonal density in ESC media 2 days after transfection. After 5-6 days, single colonies were picked and plated on 96-well plates. These plates were then duplicated 2 days later and individual clones were screened for the desired mutation by PCR followed by restriction fragment length polymorphism (RFLP) analysis. Cell lysis in 96-well plates, PCR on lysates, and restriction digests were performed. The presence of the desired *Lin28a-GFP* or *Lin28a-GFP-NLS* insertions in putative clones was confirmed by Sanger sequencing.

As C-terminally tagged GFP labeled LIN28A transgenes were shown to be able to repress let7 miRNA biogenesis (Piskounova et al., 2011) and regulate gene expression changes independent of let7 (Balzer et al., 2010), the tagging of endogenous *Lin28a* was also performed at the C-terminus. For the generation of *Lin28a-GFP* or *Lin28a-GFP-NLS* J1 ESCs, specific gRNAs targeting upstream of the stop codon (Supplementary Table) were cloned into a modified version of the SpCas9-T2A-GFP/gRNA plasmid (px458172, Addgene plasmid #48138), where we fused a truncated form of human Geminin (hGem) to SpCas9 in order to increase homology-directed repair efficiency generating SpCas9-hGem-T2A-GFP/gRNA. To generate *Lin28a-GFP* or *Lin28a-GFP-NLS* targeting donors, dsDNA gene fragments were synthesized and cloned into a vector carrying ~400 bp homology arms synthesized (IDT, Coralville, IA, USA) (Supplementary Table).For targetings, wild-type ESC were transfected with a 4:1 ratio of donor oligo and SpCas9-hGem-T2A-GFP/gRNA construct. Positively transfected cells were isolated based on GFP expression using fluorescence-activated cell sorting (FACS) and plated at clonal density in ESC media 2 days after transfection. After 5–6 days, single colonies were picked and plated on Optical bottom μClear 96-well plates and re-screened for the correct expression and localization of eGFP using live-cell spinning-disk confocal imaging. These plates were then duplicated 2 days later and individual clones were screened for the desired mutation by PCR followed by restriction fragment length polymorphism (RFLP) analysis. Cell lysis in 96-well plates, PCR on lysates, and restriction digests were performed. The presence of the desired Lin28a-GFP or *Lin28a-GFP-NLS* insertions in putative clones was confirmed by Sanger sequencing. Using the same methodological approach we generated and validated Lin28a-GFP *Npm1-HALO* ESC line by using Lin28a-GFP ESC clone to further tag HALO at C-terminus of *Npm1,* thereby generating a double KI line.

To generate *Lin28a* knockout cells, we used two independent targeting strategy allowing us to verify the effect of LIN28A KO independent of the use of individual gRNAs. The first strategy entailed the use of two gRNAs with target sites flanking the second *Lin28a* exon to excise it on both alleles. gRNA oligos were cloned into the SpCas9-T2A-PuroR/gRNA vector (px459) via cut-ligation (Supplementary Table). ESCs were transfected with an equimolar amount of each gRNA vector. Two days after transfection, cells were plated at clonal density and subjected to a transient puromycin selection (1 μg/mL) for 40 h. Colonies were picked 6 days after transfection and PCR primers were used to identify clones in which the *Lin28a* second exon had been removed. Removal of the *Lin28a* second exon was confirmed with Sanger sequencing and loss of LIN28A expression was assessed by Western Blot. To additionally corroborate the formation of multilumen embryos we generated *Lin28a* knockout cells by gRNA targeting the second exon (Supplementary Table) using the LIN28A-GFP ESC line that was used to generate morphologically normal embryos. Positively targeted LIN28A KO cells were isolated based on diminished GFP expression using fluorescence-activated cell sorting (FACS) and plated at clonal density onto Optical bottom μClear plate. After 4–5 days, colonies were screened for the diminished expression of eGFP using live-cell spinning-disk confocal imaging. Clones were subsequently genotyped and efficient *Lin28a* depletion was assessed by Western Blot.

The Sox2Δb.s mESC line was generated by deleting a ~100 nt region entailing the LIN28A binding cluster. Two gRNAs with target sites flanking the LIN28A binding cluster were used to excise it on both alleles. gRNA oligos were cloned into the SpCas9-T2A-PuroR/gRNA vector (px459) via cut-ligation (Supplementary Table). ESCs were transfected with an equimolar amount of each gRNA vector. Two days after transfection, cells were plated at clonal density and subjected to a transient puromycin selection (1 μg/mL) for 48 h. Colonies were picked 7 days after transfection and PCR primers were used to identify clones in which the *Sox2* 3’UTR sequence exon had been excised, which was additionally confirmed by Sanger sequencing.

For CRISPR/Cas gene editing, all transfections were performed using Lipofectamine 3000 (Thermo Fisher Scientific) according to the manufacturer’s instructions. All DNA oligos used for gene editing and screening are listed in Supplementary Table.

### Generation of a stable inducible LIN28A-GFP mESC line

To generate inducible PiggyBac system, the LIN28A-eGFP constructs were were synthesized as gBlocks (IDT, Coralville, IA, USA) and inserted into the enhanced PiggyBac (ePB) vector for stable integration (De Santis et al., 2018). This plasmid contains a TET-on system for inducible transgene expression. mESCs were co-transfected with 4 μg of transposable vector and 1 μg of the piggyBac transposase using Lipofectamine 3000 (Thermo Fisher Scientific) according to the manufacturer’s instructions. Selection in 5μg/ml blasticidin was initiated 3 days after transfection and maintained until resistant colonies became visible. Individual resistant inducible clones were selected, expanded and checked for LIN28A-GFP expression. 2 verified clones were used for studies including Quantseq (Fig. 4C). The LIN28A-GFP overexpression was induced by adding 1μg/ml doxycycline (dox) (ThermoFisher Scientific) in 2iLIF medium.

### Tetraploid complementation

Tetraploid chimeras were generated according to standard protocols (Engert et al., 2013; Modic et al., 2019). Embryos were collected from the mT/mG expressing mouse line (Muzumdar et al., 2007), maintained on C57/Bl6J background.

### Animal data

Mouse keeping was done at the central facilities at HMGU in accordance with the German animal welfare legislation and acknowledged guidelines of the Society of Laboratory Animals (GV-SOLAS) and of the Federation of Laboratory Animal Science Associations (FELASA).

### Immunofluorescence of embryos

Immunofluorescence whole-mount staining was performed in the following way. Briefly, embryos were isolated, fixed for 20 minutes using 2% PFA in PBS, permeabilized using 0.1% Triton X-100 in 0.1 M glycine pH 8.0. After blocking using 10% FCS, 3% donkey serum, 0.1% BSA, 0.1% Tween 20 for 2 h, embryos were incubated with the primary antibody o/n at 4C in blocking solution. After several washes in PBS containing 0.1% Tween-20 (PBST) embryos were incubated with secondary antibodies (donkey anti-goat 488, donkey anti-rabbit 555 each 1:800) in blocking solution for 3 h. During the final washes with PBST, embryos were stained with 40,6-diamidino-2-phenylindole, dihydrochloride (DAPI), transferred into 15% and 30% glycerol and embedded between two coverslips using 120m m Secure-Seal spacers (Invitrogen, S24737) and ProLong Gold antifade reagent (Invitrogen, P36930). Antibodies: Foxa2 (Santa Cruz, sc-6554), Brachyury (N-19, Santa Cruz, sc17743), RFP 1:1000 (Rockland, 600-401-379S), SOX2 (Y-17, Santa Cruz, sc-17320), Oct-3/4 (N-19, Santa Cruz, sc-8628), anti-rabbit IgG 555 (Invitrogen, A31572), donkey anti-chicken IgY (Dianova, 703-225-155), donkey anti-goat IgG 633 (Invitrogen, A21082).

### Cell culture Immunofluorescence

Cells were grown in Geltrex-coated (Thermo, A1569601) 8 well chamber slides (80826, Ibidi) and fixed with 4% PFA/DPBS solution (Thermo Fisher Scientific 16% Formaldehyde (w/v), Methanol-free, 28906) for 15 min at RT and permeabilized using 0.3% Triton X-100/DPBS solution for 15 min at RT. Primary and secondary antibodies were diluted per manufacturer recommended concentrations in 10%FBS/0.3% Triton X-100/DPBS and incubated for 1 hour at RT using the following antibodies Tbx3 (sc-17871 Santa Cruz); Nr5a2 (PP-H2325-00 R&D Systems); LIN28A (A177 Cell Signaling and ab63740 Abcam); Klf4 (AF3158, R&D Systems); Nanog (8822, Cell Signaling); Sox2 (2748S, Cell Signaling); Klf2 (Mab5466, R&D Systems); Donkey anti-rabbit IgG 555 (A31572, Invitrogen); Donkey anti-goat IgG 488 (A11055, Invitrogen); donkey anti-goat IgG 633 (Invitrogen, A21082). The samples were washed with DAPI (50ug/ml) solution and imaged using Nikon Ti2 spinning disk confocal microscope using the parameters as described in the next paragraph.

### Live cell imaging

For time lapse imaging, ESCs expressing LIN28A-GFP and NPM1-Halo were plated on Geltrex (A1569601, Gibco) coated Ibidi μ-Slide 2 Well plates (80286, Ibidi). Halo fusion-protein was labeled using with 50 nM Janelia Fluor® 646 HaloTag® for 30 minutes, then washed 4 times with PBS. Timelapse imaging was carried out on a spinning disk confocal microscope consisting of a Nikon Ti2 microscope equipped with a Yokogawa CSU-W1 spinning-disk confocal unit (50 μm pinhole size), a Prime 95B (Teledyne-Photometrics) sCMOS camera, a Nikon 60×/1.45 NA Plan Apo oil immersion objective, a motorized stage and a Perfect Focus System (Nikon). The imaging system is equipped with an environmental chamber maintained at 37 °C with 5% CO2 (Okolab). Images were acquired with the 488, 561, and 640 nm laser lines, full-frame (1200 x 1200) with 1 × 1 binning, and with a pixel size of 183 nm. The microscope was controlled by software from Nikon (NIS Elements, ver. 5.02.00). Images were acquired every 15 minutes for 13 hours at 30 different fields of view, using a 488 nm laser for Lin28-GFP, and a 650 nm laser for NPM1-Halo.

### Super-resolution cell imaging

Steady state super-resolution imaging of live ESCs and immunofluorescence images were acquired using a immunoOlympus IX83 microscope equipped with a VT-iSIM super resolution imaging system (Visitech International), Prime BSI Express scientific sCMOS camera (Teledyne-Photometrics) was used with an Olympus 150x/1.45 TIRF Apochromat oil immersion objective. The imaging system is equipped with a motorised stage with piezo Z (ASI) and environmental chamber maintained cells at 37 °C with 5% CO2 (Okolab). Images were acquired with the 405, 488, 561, and 640 nm laser lines, image size 1541 x 1066 with 1 × 1 binning, and with a pixel size of 43 nm. The microscope was controlled with μManager v2.0 gamma software (Edelstein et al.). ER-Tracker (E12353, Thermo) was used for the staining of the cytoplasm in live cells. Deconvolution was done using Huygens (SVI).

### Imaging of the embryogenesis system

Images were acquired using a Leica TCS SP8 DLS AOBS confocal microscope equipped with a DMi8 microscope stand with Galvo Z stage, adaptive focus control, a 1x scCMOS camera (Leica DFC9000 GTC DLS, 2048 x 2048 pixels), 2x spectral PMT and 2x spectral HyD detectors, 40x/1.30NA immersion oil HC PL APO CS2 and 63x/1.40NA immersion oil HC PL APO CS2 objectives. Images were acquired using 405, 488, 561, 594 and 633nm lasers, image size 1024×1024. The microscope was controlled with LAS-X software.

### Cytoplasm and nucleus intensity analysis

For quantitative analysis of Lin28-GFP levels, we used Cellpose (Stringer et al., 2021) to carry out segmentation of nuclei in the NPM1-HALO channel from images at all timepoints and fields of view in the time lapse imaging. Masks generated with Cellpose were then filtered in Fiji for size and roundness to discard small round dead cells and rounded mitotic cells from analysis, and objects on the edges of the image were removed. Remaining masks were then used to measure intensity of Lin28A inside the nucleus. To measure intensity in the cytoplasm, we enlarged the masks by 3 pixels, and took the intensity along the perimeter of the enlarged mask as a readout for intensity in the cytoplasm (discarding pixels which lay in neighbouring masks), similarly to cytoplasm intensity measurements carried out elsewhere (Mulholland et al., 2020).

### Flow cytometry

For flow cytometry experiments, single-cell suspensions were made using Accutase (A6964, Sigma-Aldrich) for 5 min in 37 °C; or Enzyme-Free Cell Dissociation Buffer (13151014, Gibco) for 30 min at 37 °C and washed with 5% FBS (EmbryoMax® ES Cell Qualified FBS, ES-009-B, Merck) in PBS, incubated with fluorophore-conjugated antibodies against SSEA1 (MC480, Thermo) or SSEA4 (MC813-70, Thermo) for 30-60 min on ice. Cells were centrifuged, resuspended in buffer containing SYTOX blue for dead cell exclusion and analyzed using a FACS Aria III (BD Biosciences). Cell debris were excluded by forward and side scatter gating. FlowJo was used for data analysis.

For intracellular flow cytometry mESCs were dissociated by Accutase (A6964, Sigma-Aldrich), centrifuged and resuspended in 2% of methanol-free formaldehyde (28906, Thermo Fisher Scientific) for 10 min in RT. Inside Stain kit (130-090-477, Miltenyi Biotec) was used according to manufacturer’s protocol using Alexa647-conjugated SOX2 antibody (D6D9-5067, Cell Signaling).

### Western blotting

Cells were trypsinized and lysed using RIPA buffer, containing phosphatase (Sigma-Aldrich, 4906837001) and protease (Merck, 539134) inhibitors. After addition of 4x LDS loading buffer (Thermo, NP0007) with 2-Mercaptoethanol (Sigma-Aldrich, M3148) samples were heated to 95Cfor 5 min. Samples were ran on NuPAGE™ 4 to 12%, Bis-Tris (Thermo, NP0321PK2), and blotted using the Power Blotter XL System (Thermo, PB0013). Following 3x 5 min washes with PBS-T, membranes were blocked with 5% milk powder (T145.1, Carl Roth) in TBS-T. Membranes were then incubated o/n at 4C with 5% milk powder in PBS-T containing the primary antibodies at the following dilutions (1:1000 Lin28a (A177, Cell signaling); 1:4000 H3 (Abcam, ab1791), GAPDH (Cell Signaling, 2118S)). After 3x 5 min PBS-T washes, membrane was incubated with goat anti-rabbit IgM-HRP (sc-2030, Santa Cruz) or Goat anti-mouse IgG HRP-conjugated (Dianova, 115-035-003) in 5%milk powder in PBS-T. Following 4x 10 min washes with PBS-T the membrane was incubated for 1 min with Clarity Western ECL Substrate (170-5060, Bio-Rad Laboratories) and imaged with Amersham™ Imager 600 (GE, 29-0834-61).

### Cell fractionation

~10^7 cells were resuspended in 500μl of buffer S (10 mM HEPES, [pH 7.9], 10 mM KCl, 1.5 mM MgCl 2, 0.34 M sucrose, 10% glycerol, 0.1% Triton X-100, 1 mM DTT, proteinase inhibitor (04693116001, Roche). The cells were incubated for 5 min on ice. Nuclei were collected into a pellet by low-speed centrifugation (4 min, 1,300 × g) at 4°C. The supernatant containing cytoplasm fraction was further clarified by high-speed centrifugation (15 min, 20,000 × g, 4°C) to remove cell debris and insoluble aggregates. To further purify the nuclei, these were washed three times in buffer S and spun down (1700g for 5min) at 4°C to get soluble nuclear fraction.

### mRNA-RBP occupancy (mRNA interactome capture)

Protocol was performed as described in (Modic et al., 2019). 3 biological replicates of ~10^7 nESC, RSC, EpiSC were used and the comparable amount of T+ mesodermal progenitors was FACS sorted in 3 replicates. Different developmental stages of ESC were irradiated as described for iCLIP, apart from non-crosslinked mock oligo(dT) pull-downs. To further strengthen a cohort of *bona fide* mRNA-RBP occupancy, apart from non-crosslinked mock oligo(dT) pull-downs, the control naked beads lacking oligo(dT) wew implemented to eliminate not only proteins that unspecifically bind oligo(dT) but also common contaminants of magnetic beads

### FASP digest

Each 10mg of RBPome were digested with a modified FASP procedure (Wiśniewski et al., 2009). Briefly, the proteins were reduced and alkylated using dithiothreitol and iodoacetamide, diluted with one volume of UA buffer (8Murea in 0.1M Tris/HCl pH 8.5) and then centrifuged through a 30 kDa cut-off filter device (PALL, Port Washington). Samples were washed twice with UA buffer and twice with 50mM ammonium bicarbonate prior to digest of the immobilized proteins on the filter for 2 h at RT using 1 mg Lys-C (Wako Chemicals) and for 16 h at 37C using 2 mg trypsin (Promega). Tryptic peptides were collected by centrifugation (10 min at 14,000 g), and the samples were acidified with 0.5% TFA and stored at 20C.

### Mass spectrometry

Before loading, the samples were centrifuged for 5 min at 4C. LC-MS/MS analysis was performed on a QExactive HF mass spectrometer (Thermo Scientific) online coupled to an Ultimate 3000 nano-RSLC (Thermo Scientific). Approximately 0.5 mg of every digested sample was automatically injected and loaded onto the trap column at a flow rate of 30 ml/min in 3% ACN/ 0.1% FA. After 5 min, the peptides were eluted from the trap column and separated on the C18 analytical column (75 mm i.d. x 25 cm, Acclaim PepMap100 C18.2 mm, 100A°, Dionex) by a 90 min gradient from 5 to 25% ACN in 0.1% FA at 300 ml/min flow rate followed by a 5 min gradient from 25% to 40% ACN in 0.1% FA. Between each sample, the column was washed with 85% ACN for 5 min followed by equilibration at 3% ACN in 0.1% FA for 18 min. MS spectra were recorded at a resolution of 60000 with an AGC target of 3e6 and a maximum injection time of 50 ms from 300 to 1500 m/z. From the MS scan, the 10 most abundant peptide ions were selected for fragmentation via HCD with a normalized collision energy of 27, an isolation window of 1.6 m/z and a dynamic exclusion of 30 s. MS/MS spectra were recorded at a resolution of 15000 with a AGC target of 1e5 and a maximum injection time of 50 ms. Intensity threshold was set to 1e4and unassigned charges and charges of +1 and > 8 were excluded.

### LC-MS/MS analysis of the total proteome and mRNA-RBP interactome (RBPome)

Raw files from total proteome (nESC, RSC, EpiSC and T+ mesodermal progenitors, 3 replicates each) and RNA-interactome (nESC, RSC, EpiSC and T+ mesodermal progenitors, 3 replicates each, and naked-beads (5 replicates, 1 from nESCs and 2 from RSCs and EpiSCs) and no-UV controls (3 replicates, 1 from nESCs, RSCs and EpiSCs)) were processed together with MaxQuant version 1.6.6.0 as separate parameter groups, using separate label free quantification, enabled match between runs within parameter groups and disabled large LFQ ratio stabilization as non-default options. Spectra were searched against forward and reverse sequences of the mouse proteome (ensembl, GRCm38 release 96). Carbamidometylation was set as fixed, and methionine oxidation and N-acetylation as variable modification. R version 4.0.2 was used for downstream analysis. Naked-beads and no-UV samples were grouped as control samples. Protein groups identified as contaminants, on basis of razor or reverse peptides were excluded from analysis.

LFQ values were log2 transformed and two-stage normalisation was performed on the basis of proteins, quantified in all samples within sample groups (first stage within RBPomes, protemes and controls, second stage within both RBPomes and controls). Missing value imputation, with values randomly sampled from normal distribution shrunk by a factor of 0.3 and downshifted by 1.8 standard deviations, was performed for RBPome and control, and proteome samples separately. Uniform Manifold Approximation and Projection (UMAP) dimensionality reduction was performed using R package umap version 0.2.6.0. Only protein groups identified in all replicates of RBPome and proteome of at least one cell state were considered for analysis of mRNA-interactome and dynamic binding (976). Only protein groups identified in all replicates of proteomes of at least one cell state were considered for proteome analysis (3876).

In order to evaluate cross-linking efficiency of individual samples from different cell states, a pre-analysis of enrichment of protein groups in RBPomes over controls was performed using generalized linear model (R base package *stats)* with model matrix expressed in R notation as *log2(LFQ) ~ cell_state,* where *cell_state* is a designation for control (intercept), naïve ESCs, RSCs, EpiSCs or T+ RBPome samples. Protein groups with thus obtained gaussian log2 fold enrichment > 2 and p-value < 0.01 across all conditions were used for enrichment effect scaling. Median enrichments were calculated for individual replicates of all cell states over controls and divided by their median. Thus calculated effect scales were close to 1 (meaning no correction and highly similar cross-linking efficiency), ranging from 0.94 – 1.03, and were used in further analysis where applicable. For all further analysis-proper, protein-wise lasso-based linear models (R package *glmnet,* version 4.0.2, (Friedman et al., 2010)) were used for their stringency and sparsity inducing property, leveraging fixed lasso inference (R package *selectiveInference,* version 1.2.5, (Tibshirani et al., 2014)) for p-value estimation. In using package *glmnet,* specifically function *cv.glmnet,* default parameters were used with exception of: *thresh* was set to 1E-28, *maxit* was set to 1E7 and *nfolds* was set to 10. In using package *selectiveInference,* specifically function *fixedLassoInf,* default parameters were used with exception of: models at *lambda(min)* were used except in analysis of dynamic binding where *lambda(min* + *1se)* was used for increased stringency towards estimating effect interactions, *tol.beta* was set to 0.01, *tol.kkt* was set to 0.01, *sigma* was explicitly calculated using built-in function and number of *bits* used for numeric calculation was set to be minimum of 100 and increased if convergence was not reached. If numeric convergence was not reached using 500 *bits,* the protein group was dropped from the analysis.

Enrichment of protein groups in RBPomes over controls was obtained from protein-wise fitted lasso-based linear model with model matrix expressed in R notation as *log2(LFQ) ~ cell_state,* where *cell_state* is a designation for control (intercept), naïve ESCs, RSCs, EpiSCs or T+ RBPome samples. Effects and p-values were calculated at *lambda(min).* Protein groups with log2 fold-enrichment above 4 and p-value below 0.00001 in all sample groups were considered bona-fide RBPs and used for further analysis (284, with 365, 359, 376, 359 and 464 enriched in naïve, LIMES, primed, T+ or any of the cell states, respectively). Non-reduntant GO-term mapping to each protein group was performed using R package biomaRt version 2.45.5. GO-term analysis was performed using fisher exact test with fdr-based p-value adjustment. List of bona-fide mouse transcription factors related to Supplementary Figure 5E was taken from RIKEN TFdb (Kanamori et al., 2004).

Significant deviation of change in total proteome intensity and RNA-interactome intensity between cell states (dynamic binding) was determined using model matrix expressed in R notation as *log2(LFQ) ~ dataset* + *cell_state* + *cell_state:dataset,* where *dataset* represents Proteome or RBPome and *cell_state* represents naïve ESCs, RSCs, EpiSCs or T+ cell states. Effects and p-values were calculated at *lambda(min+1se).*The effects describing interaction of *cell_state* and *dataset* (in R notation *cell_state:dataset)* were taken as a measure of differential RNA binding, with proteins for which this effect was in absolute manner larger than 0.25 and p value below 0.001 considered dynamic RNA binding proteins. In total, 9, 14 and 48 bona-fide RNA binding proteins were considered significant dynamic binders in comparison of RSCs, EpiSCs and mesoderm progenitors with naive state, respectively.

Analysis of changes in the full proteome and RBPome was performed in similar manner, using model matrix expressed in R notation as *log2(LFQ) ~ cell_state,* where cell_state is a designation for naïve ESCs (intercept), RSCs, EpiSCs or T+ proteome or RBPome samples.

Effects and p-values were calculated at *lambda(min).* Protein abundance changes, in absolute manner larger than 1 (log2) with associated p value below 0.0001 were considered significant. In mRNA interactome analysis, 3, 10 and 18 bona-fide RNA binding proteins were considered significantly changed in comparison of RSCs, EpiSCs and mesoderm progenitors with naive state, respectively. In full proteome analysis, 109, 479 and 686 proteins were considered significantly changed in comparison of RSCs, EpiSCs and mesoderm progenitors with naive state, respectively.

### RNA preparation, qRT-PCR assays and RNA sequencing

Total RNA was prepared from cell pellets using miRNeasy Micro Kit (Qiagen, 217084) according to the manufacturer’s instructions. miR-let7 levels were analyzed using the miScript protocol (Qiagen, 218193) using primers amplifying miR let7 (Qiagen, MS00005866; let-7g Qiagen, MS00010983).

For chimera RNA sequencing we first FACS sorted GFP-expressing 500 cells and used microcapillary electrophoresis on Agilent 2100 Bioanalyzer with RNA Pico 6000 kit (Agilent, 5067-1513) to analyze RNA quality (RIN values > 8). For total RNA sequencing Ovation RNA Amplification System V2 was used according to manufacturer’s instruction (Nugen, 3100-A01). For 3’mRNA-seq, 0.5 μg of totalRNA was used per library prepared using Lexogen QuantSeq-FWD kit (Lexogen GmbH, 0.15) according to manufacturer’s instructions. All libraries were evaluated on an Agilent 2200 TapeStation using the High Sensitivity D1000 ScreenTape (Agilent, 5067-5585) and DNA concentration was measured using a promega Quantifluor® dsdna system (Promega, E2670). Samples were sequenced using HiSeq4000 to generate 100-nt single-end reads.

### SLAMseq

Thiol(SH)-linked alkylation for the metabolic sequencing of RNA (SLAMseq) has been generated according to the published protocol (Ameres et al., 2017). Naïve or priming ESC were treated with 100 μM 4-thiouridine (s4U) (Sigma, T4509-25MG) for 24hours of pulse labeling. During the last 15hours of pulse labeling, fresh s4U-containing media has been exchanged every 3hours to enhance s4U incorporation in RNA species. Prior to initiation of the chase labeling with 10mM Uridine-containing (Sigma, U3750-25G) growth medium (100X excess of uridine compared to initial s4U concentrations), the media from the cells have been removed and washed twice with chase-labeling medium. The chase-labeling was initiated upon washing off pulse-labeling medium and upon indicated time-points, the media was taken off and cells were lysed directly in Qiazol (Qiagen, 79306), followed by RNA preparation.

### RNA-seq processing and analysis

Reads were quantified using Salmon (Patro et al., 2017) for iLIN28A-GFP overexpression analysis or using STAR (Dobin et al., 2013), with a transcriptome built from GENCODE M22 transcripts. Additionally, to monitor the expression level of the Lin28a-GFP transgene, the sequence was included with decoy sequences derived from the whole genome sequence. Differential expression analysis was performed using DESeq2 (Love et al., 2014), with effect size shrinkage using the apeglm package (Zhu et al., 2019). For iLIN28A-GFP overexpression analysis, the developmentally down-regulated genes were defined as genes at least two-fold down-regulated at the 24 hour and 48 hour timepoints in previously published data (Yang et al., 2019b), and a list of naive pluripotency genes was derived from the same study.

### RNA stability prediction methods

To calculate mRNA T>C conversion the pipeline was enclosed in Snakemake5.3.0 (Koster and Rahmann, 2012). For analysis of mRNA 3’ end sequencing data, reads were demultiplexed with Cutadapt 1.18 prohibiting mismatches in the barcode (Martin, 2011). Then, sequencing reads quality was assessed (-q 20), barcode trimmed (--cut 12) discarding short reads (--length 16). Poly-adenylated stretches (>4) at the 3’ end were removed up to 20 tandems without indels nor mismatches. Trimmed reads were aligned with Slamdunk map0.3.3 (Neumann et al., 2019) to reference mouse genome (mm10) using local alignment and allowing up to 100 multi-mapping reads (-n 100). Aligned reads were subject to a filtering step retaining only alignments with a minimum identity of 95% and a minimum of 50% of the read bases mapped. Reads ambiguously mapping to more than one 3’UTR were discarded (https://github.com/AmeresLab/UTRannotation). SNPs were called using VarScan 2.4.1with default parameter and establishing 10 fold coverage cut off and a variant fraction cut off 0.8 (-c 10 -f 0.8). SNPs overlapping T> C conversions need to present at least >26 base quality Phred score. T> C conversions were counted with rolling windows and normalised by T content, coverage of each position and average per possible transcript 3’UTR. Finally, the T> C conversion quality was assessed by Slamdunk alleyoop0.3.3 (Neumann et al., 2019).

To calculate half-life, T> C conversion were background-subtracted and normalised to the chase S4U treatment. T> C conversions were fit to exponential decay non-linear function using the R minpack.lm 1.2 (Bates and Chambers, 2017). Only 3’UTR half-life that correlated to the model R2 > 0.6 in all the experimental conditions were retrained.

### Pre-processing and feature computation or model training

Initially, 3’UTRs were filtered for R^2 > 0.6 and a delta half life was computed (Naive – Primed and Lin28 KO – LIN28 WT). All retained 3’UTRs were sorted according to their delta half life. Top and bottom 1000 3’UTRs with the largest half life changes were assigned to a specific condition e.g. Naive and Primed.

Next, we computed a feature matrix for these 3’UTR. Features were distinguished into seven classes and computed gbm as described below:

| Feature class | Feature description | #Features |
| --- | --- | --- |
| Simple features | GC content, Dinucleotide frequency, 3'UTR width, average PhastCon score | 19 |
| miCLIP-seq | Average m6A miCLIP-seq scores measured in different ESC conditions | 6 |
| ESC iCLIP-seq | Average CLIP-seq scores for various RBPs measured in mouse ESCs | 62 |
| TAIL-seq [REF] | poly(A) length, #A, #monoC, #monoG, #monoT, #oligoC, #oligoG, #oligoT from mouse ESCs | 8 |
| TargetSCAN [REF] | Binding predictions for different miRNAs | 11 |
| DeepBind [REF] | Average binding prediction of 50bp tiled 3'UTRs | 105 |
| POSTAR2 [REF] | CLIP-seq peak scores from various RBPs and conditions | 54 |

All these features were assembled in one matrix, filtered for zero variance features and split into a training set (75%) and a hold out test set (25%). The training set was used for subsequent model training (one model per set and one model for all features together).

For Figure 3 F, we compared the univariate feature importance, as measured by the auROC for each CLIP/Postar2 feature. auROCs were computed per feature, once to distinguish WT vs KO and Naive vs Primed 3’UTRs. For this analysis we used the varImp function from the R-package caret.

### Gradient boosting machine (GBM) training and model benchmarking

Hyperparameter search and model training was performed with the R-package caret as well as the GBM package (Greenwell et al., 2019). Models were trained for a binary classification task to distinguish LIN28 WT against KO or respective Naive against Primed specific 3’UTRs. Model training was performed with a 5 times repeated 5 fold cross validation approach using the training set as input. Further, the hyperparameter search was performed as grid search over the parameters listed in the table below with the aim to maximise the auROC.

| Parameter | Value |
| --- | --- |
| Interaction depth | 1, 3, 6 |
| Number of trees | 1, 10, 20, 30, 40, 50, 100, 150, 200, 250, 300, 350, 400, 450, 500 |
| Shrinkage | 0.001, 0.005, 0.01, 0.05, 0.1 |
| Min number of observations in terminal node | 10 |

Resulting GBM models were evaluated on the hold-out test set using the auROC as well as Matthews correlation coefficient (MCC) as performance measures.

Feature importance for each GBM model were computed as relative influences as well as reduction in model performance by shuffling a particular feature. For this analysis we used the associated GBM R package function “relative.influence” and “permutation.test.gbm”.

### iCLIP

We used iCLIP to identify a repertoir of RBP binding sites in mESCs. Cells were UV crosslinked on ice and then lysed in RIPA buffer. 0.4 Units of RNaseI and 4 Units Turbo DNase were added per 1 ml of cell lysate at 1mg/ml protein concentration for RNA fragmentation. Negative controls (no-UV) were prepared. Antibodies were coupled to magnetic Protein G beads used to isolate Protein-RNA complexes, and RNA was ligated to a pre-adenylated infra-red labelled IRL3 adaptor (Zarnegar et al., 2016) with the following sequence:

*/5rApp/AG ATC GGA AGA GCG GTT CAG AAA AAA AAA AAA /iAzideN/AA AAA AAA AAA A/3Bio/*

The complexes were then size-separated by SDS-PAGE, blotted onto nitrocellulose and visualised by Odyssey scanning. For the multiplexed sample 1 replicated was run in parallel with the IRL3 to allow quality control of the RNP complex on the membrane and to help with cutting of the bands. RNA was released from membrane by proteinase K digestion and recovered by precipitation. cDNA was synthesized with Superscript IV Reverse Transcriptase (Life Technologies) and AMPure XP beads purification (Beckman-Coulter, USA), then circularised using Circligase II (Epicentre) followed by AMPure XP beads purification. After PCR amplification, libraries were size-selected with Ampure beads (if necessary by gel-purification) and quality controlled for sequencing. Libraries were sequenced as single end 100bp reads on Illumina HiSeq 4000.

### Visualisation of iCLIP data

For the comparative visualisation of iCLIP data, first we normalised iCLIP cDNA/crosslink counts at a given crosslink site by the experimental library size. Then, normalised counts were Smoothed with a gaussian kernel of width 9. The smoothed, normalised values were plotted across the region.

### iCLIP data analysis

iCLIP reads were analysed on the iMaps server (https://imaps.genialis.com/) using the iCount software (https://github.com/tomazc/iCount). Briefly, experimental barcodes were removed (Martin, 2011) and sequencing reads aligned with STAR (Dobin et al., 2013) to mouse genome build (GRCm38.p5 GENCODE version 15 annotation) allowing two mismatches. Unique Molecular Identifiers (UMIs), were used to distinguish and remove the PCR duplicates. To determine protein-RNA contact sites, the sequencing read preceding nucleotide was assigned as the cross-link event. Significant contact sites were then identified, using the iCount function, based on false discovery rate (FDR) <0.05 comparing specific sites within a window of three nucleotides with randomised data (100 permutations) within co-transcribed regions. The significant cross-links signal was normalised by sequencing deep and million of tags (CPM).

Then replicates were merged and summary of cDNA counts within genes and genic regions were generated with iCount segments function, normalising the counts by the length of the corresponding region.

### Statistics and reproducibility

No statistical methods were used to predetermine sample size, the experiments were not randomized. For microscopy analysis, where possible, experimenter bias was avoided by selecting fields of view (or individual cells) for acquisition of LIN28A-GFP signal using the NPM1 nuclear signal. To further reduce bias, imaging analysis was subsequently performed indiscriminately on all acquired images using semi-automated analysis pipelines using Fiji scripts. All the experimental findings were reliably reproduced in independent experiments as indicated in the Figure legends. In general, all micrographs from immunofluorescence and immunoblots depicted in this study are representative of *n* ≥ 2 independent experiments. The number of replicates used in each experiment are described in the figure legends and/or in the Methods section, as are the Statistical tests used including the adjustment methods for adjusted *P* values are given.

## REFERENCES

1. M. N. Shahbazi, A. Scialdone, N. Skorupska, A. Weberling, G. Recher, M. Zhu, A. Jedrusik, L. G. Devito, L. Noli, I. C. Macaulay, C. Buecker, Y. Khalaf, D. Ilic, T. Voet, J. C. Marioni, M. Zernicka-Goetz, Pluripotent state transitions coordinate morphogenesis in mouse and human embryos. Nature. 552, 239–243 (2017).

2. I. Bedzhov, M. Zernicka-Goetz, Self-organizing properties of mouse pluripotent cells initiate morphogenesis upon implantation. Cell. 156, 1032–1044 (2014).

3. A. Neagu, E. Genderen, I. Escudero, L. Verwegen, D. Kurek, J. Lehmann, J. Stel, R. A. M. Dirks, G. Mierlo, A. Maas, C. Eleveld, Y. Ge, A. T. Dekker, R. W. W. Brouwer, W. F. J. IJcken, M. Modic, M. Drukker, J. H. Jansen, N. C. Rivron, E. B. Baart, H. Marks, D. Berge, In vitro capture and characterization of embryonic rosette-stage pluripotency between naive and primed states. Nat. Cell Biol. 22, 534–545 (2020).

4. R. Fan, Y. S. Kim, J. Wu, R. Chen, D. Zeuschner, K. Mildner, K. Adachi, G. Wu, S. Galatidou, J. Li, H. R. Schöler, S. A. Leidel, I. Bedzhov, Wnt/Beta-catenin/Esrrb signalling controls the tissue-scale reorganization and maintenance of the pluripotent lineage during murine embryonic diapause. Nat. Commun. 11, 5499 (2020).

5. M. Li, J. C. I. Belmonte, Deconstructing the pluripotency gene regulatory network. Nat. Cell Biol. 20, 382–392 (2018).

6. L. Weinberger, M. Ayyash, N. Novershtern, J. H. Hanna, Dynamic stem cell states: naive to primed pluripotency in rodents and humans. Nat. Rev. Mol. Cell Biol. 17, 155–169 (2016).

7. P. Yang, S. J. Humphrey, S. Cinghu, R. Pathania, A. J. Oldfield, D. Kumar, D. Perera, J. Y H. Yang, D. E. James, M. Mann, R. Jothi, Multi-omic Profiling Reveals Dynamics of the Phased Progression of Pluripotency. Cell Systems. 8, 427–445 (2019).

8. V. A. Herzog, B. Reichholf, T. Neumann, P. Rescheneder, P. Bhat, T. R. Burkard, W. Wlotzka, A. von Haeseler, J. Zuber, S. L. Ameres, Thiol-linked alkylation of RNA to assess expression dynamics. Nat. Methods. 14, 1198–1204 (2017).

9. H. Chang, J. Lim, M. Ha, V. N. Kim, TAIL-seq: genome-wide determination of poly(A) tail length and 3’ end modifications. Mol. Cell. 53, 1044–1052 (2014).

10. Y. Zhu, G. Xu, Y. T. Yang, Z. Xu, X. Chen, B. Shi, D. Xie, Z. J. Lu, P. Wang, POSTAR2: deciphering the post-transcriptional regulatory logics. Nucleic Acids Res. 47, 203–211 (2019).

11. B. Alipanahi, A. Delong, M. T. Weirauch, B. J. Frey, Predicting the sequence specificities of DNA- and RNA-binding proteins by deep learning. Nat. Biotechnol. 33, 831–838 (2015).

12. S. C. Kwon, H. Yi, K. Eichelbaum, S. Föhr, B. Fischer, K. T. You, A. Castello, J. Krijgsveld, M. W. Hentze, V. N. Kim, The RNA-binding protein repertoire of embryonic stem cells. Nat. Struct. Mol. Biol. 20, 1122–1130 (2013).

13. A. G. Baltz, M. Munschauer, B. Schwanhäusser, A. Vasile, Y Murakawa, M. Schueler, N. Youngs, D. Penfold-Brown, K. Drew, M. Milek, E. Wyler, R. Bonneau, M. Selbach, C. Dieterich, M. Landthaler, The mRNA-bound proteome and its global occupancy profile on protein-coding transcripts. Mol. Cell. 46, 674–690 (2012).

14. D. Kurek, A. Neagu, M. Tastemel, N. Tüysüz, J. Lehmann, H. J. G. van de Werken, S. Philipsen, R. van der Linden, A. Maas, W. F. J. van IJcken, M. Drukker, D. ten Berge, Endogenous WNT Signals Mediate BMP-Induced and Spontaneous Differentiation of Epiblast Stem Cells and Human Embryonic Stem Cells. Stem Cell Reports. 4, 114–128 (2015).

15. M. Gouti, A. Tsakiridis, F. J. Wymeersch, Y. Huang, J. Kleinjung, V. Wilson, J. Briscoe, In vitro generation of neuromesodermal progenitors reveals distinct roles for wnt signalling in the specification of spinal cord and paraxial mesoderm identity. PLoS Biol. 12, e1001937 (2014).

16. M. Modic, M. Grosch, G. Rot, S. Schirge, T. Lepko, T. Yamazaki, F. C. Y. Lee, E. Rusha, D. Shaposhnikov, M. Palo, J. Merl-Pham, D. Cacchiarelli, B. Rogelj, S. M. Hauck, C. von Mering, A. Meissner, H. Lickert, T. Hirose, J. Ule, M. Drukker, Cross-Regulation between TDP-43 and Paraspeckles Promotes Pluripotency-Differentiation Transition. Mol. Cell. 74, 951–965 (2019).

17. E. J. Vogt, M. Meglicki, K. I. Hartung, E. Borsuk, R. Behr, Importance of the pluripotency factor LIN28 in the mammalian nucleolus during early embryonic development. Development. 139, 4514–4523 (2012).

18. I. Heo, C. Joo, Y.-K. Kim, M. Ha, M.-J. Yoon, J. Cho, K.-H. Yeom, J. Han, V. N. Kim, TUT4 in concert with Lin28 suppresses microRNA biogenesis through pre-microRNA uridylation. Cell. 138, 696–708 (2009).

19. S. Lin, R. I. Gregory, MicroRNA biogenesis pathways in cancer. Nature Reviews Cancer. 15, 321–333 (2015).

20. E. Balzer, C. Heine, Q. Jiang, V. M. Lee, E. G. Moss, LIN28 alters cell fate succession and acts independently of the let-7 microRNA during neurogliogenesis in vitro. Development. 137, 891–900 (2010).

21. L. C. Orietti, V. S. Rosa, F. Antonica, C. Kyprianou, W. Mansfield, H. Marques-Souza, M. N. Shahbazi, M. Zernicka-Goetz, Embryo Size Regulates the Timing and Mechanism of Pluripotent Tissue Morphogenesis. Stem Cell Reports (2020), doi:10.1016/j.stemcr.2020.09.004.

22. G. S. Eakin, A.-K. Hadjantonakis, Production of chimeras by aggregation of embryonic stem cells with diploid or tetraploid mouse embryos. Nat. Protoc. 1, 1145–1153 (2006).

23. M. N. Shahbazi, Mechanisms of human embryo development: from cell fate to tissue shape and back. Development. 147 (2020).

24. B. Vadla, K. Kemper, J. Alaimo, C. Heine, E. G. Moss, lin-28 controls the succession of cell fate choices via two distinct activities. PLoS Genet. 8, e1002588 (2012).

25. N. Shyh-Chang, H. Zhu, T. Yvanka de Soysa, G. Shinoda, M. T. Seligson, K. M. Tsanov, L. Nguyen, J. M. Asara, L. C. Cantley, G. Q. Daley, Lin28 enhances tissue repair by reprogramming cellular metabolism. Cell. 155, 778–792 (2013).

26. X. Zhou, G. G. Nair, H. A. Russ, C. D. Belair, M.-L. Li, M. Shveygert, M. Hebrok, R. Blelloch, LIN28B Impairs the Transition of hESC-Derived β Cells from the Juvenile to Adult State. Stem Cell Reports. 14, 9–20 (2020).

27. J. Zhang, S. Ratanasirintrawoot, S. Chandrasekaran, Z. Wu, S. B. Ficarro, C. Yu, C. A. Ross, D. Cacchiarelli, Q. Xia, M. Seligson, G. Shinoda, W. Xie, P. Cahan, L. Wang, S.-C. Ng, S. Tintara, C. Trapnell, T. Onder, Y.-H. Loh, T. Mikkelsen, P. Sliz, M. A. Teitell, J. M. Asara, J. A. Marto, H. Li, J. J. Collins, G. Q. Daley, LIN28 Regulates Stem Cell Metabolism and Conversion to Primed Pluripotency. Cell Stem Cell. 19, 66–80 (2016).

## References for Materials & Methods

Ameres, S., Herzog, V.A., Reichholf, B., Ameres, S., 2017. Thiol-linked alkylation for the metabolic sequencing of RNA (SLAMseq). Protoc. Exch. https://doi.org/10.1038/protex.2017.105

Balzer, E., Heine, C., Jiang, Q., Lee, V.M., Moss, E.G., 2010. LIN28 alters cell fate succession and acts independently of the let-7 microRNA during neurogliogenesis in vitro. Development 137, 891–900.

Bates, D.M., Chambers, J.M., 2017. Nonlinear Models. Statistical Models in S. https://doi.org/10.1201/9780203738535-10

Burtscher, I., Lickert, H., 2009. Foxa2 regulates polarity and epithelialization in the endoderm germ layer of the mouse embryo. Development 136, 1029–1038.

De Santis, R., Garone, M.G., Pagani, F., de Turris, V., Di Angelantonio, S., Rosa, A., 2018. Direct conversion of human pluripotent stem cells into cranial motor neurons using a piggyBac vector. Stem Cell Res. 29, 189–196.

Dobin, A., Davis, C.A., Schlesinger, F., Drenkow, J., Zaleski, C., Jha, S., Batut, P., Chaisson, M., Gingeras, T.R., 2013. STAR: ultrafast universal RNA-seq aligner. Bioinformatics 29, 15–21.

Engert, S., Burtscher, I., Kalali, B., Gerhard, M., Lickert, H., 2013. The Sox17CreERT2 knock-in mouse line displays spatiotemporal activation of Cre recombinase in distinct Sox17 lineage progenitors: Sox17CreERT2Mouse Line. Genesis 51, 793–802.

Friedman, J., Hastie, T., Tibshirani, R., 2010. Regularization Paths for Generalized Linear Models via Coordinate Descent. J. Stat. Softw. 33, 1–22.

Greenwell, B., Boehmke, B., Cunningham, J., Developers, G., 2019. gbm: Generalized boosted regression models. R package version 2.

Kanamori, M., Konno, H., Osato, N., Kawai, J., Hayashizaki, Y., Suzuki, H., 2004. A genome-wide and nonredundant mouse transcription factor database. Biochem. Biophys. Res. Commun. 322, 787–793.

Koster, J., Rahmann, S., 2012. Snakemake--a scalable bioinformatics workflow engine. Bioinformatics. https://doi.org/10.1093/bioinformatics/bts480

Love, M.I., Huber, W., Anders, S., 2014. Moderated estimation of fold change and dispersion for RNA-seq data with DESeq2. Genome Biol. 15, 550.

Martin, M., 2011. Cutadapt removes adapter sequences from high-throughput sequencing reads. EMBnet.journal 17, 10–12.

Modic, M., Grosch, M., Rot, G., Schirge, S., Lepko, T., Yamazaki, T., Lee, F.C.Y., Rusha, E., Shaposhnikov, D., Palo, M., Merl-Pham, J., Cacchiarelli, D., Rogelj, B., Hauck, S.M., von Mering, C., Meissner, A., Lickert, H., Hirose, T., Ule, J., Drukker, M., 2019. Cross-Regulation between TDP-43 and Paraspeckles Promotes Pluripotency-Differentiation Transition. Mol. Cell 74, 951–965.e13.

Mulholland, C.B., Nishiyama, A., Ryan, J., Nakamura, R., Yiğit, M., Glück, I.M., Trummer, C., Qin, W., Bartoschek, M.D., Traube, F.R., Parsa, E., Ugur, E., Modic, M., Acharya, A., Stolz, P., Ziegenhain, C., Wierer, M., Enard, W., Carell, T., Lamb, D.C., Takeda, H., Nakanishi, M., Bultmann, S., Leonhardt, H., 2020. Recent evolution of a TET-controlled and DPPA3/STELLA-driven pathway of passive DNA demethylation in mammals. Nat. Commun. 11, 5972.

Muzumdar, M.D., Tasic, B., Miyamichi, K., Li, L., Luo, L., 2007. A global double-fluorescent Cre reporter mouse. Genesis 45, 593–605.

Neagu, A., van Genderen, E., Escudero, I., Verwegen, L., Kurek, D., Lehmann, J., Stel, J., Dirks, R.A.M., van Mierlo, G., Maas, A., Eleveld, C., Ge, Y., den Dekker, A.T., Brouwer, R.W.W., van IJcken, W.F.J., Modic, M., Drukker, M., Jansen, J.H., Rivron, N.C., Baart, E.B., Marks, H., Ten Berge, D., 2020. In vitro capture and characterization of embryonic rosette-stage pluripotency between naive and primed states. Nat. Cell Biol. 22, 534–545.

Neumann, T., Herzog, V.A., Muhar, M., von Haeseler, A., Zuber, J., Ameres, S.L., Rescheneder, P., 2019. Quantification of experimentally induced nucleotide conversions in high-throughput sequencing datasets. BMC Bioinformatics 20, 258.

Patro, R., Duggal, G., Love, M.I., Irizarry, R.A., Kingsford, C., 2017. Salmon provides fast and bias-aware quantification of transcript expression. Nat. Methods 14, 417–419.

Piskounova, E., Polytarchou, C., Thornton, J.E., LaPierre, R.J., Pothoulakis, C., Hagan, J.P., Iliopoulos, D., Gregory, R.I., 2011. Lin28A and Lin28B inhibit let-7 microRNA biogenesis by distinct mechanisms. Cell 147, 1066–1079.

Stringer, C., Wang, T., Michaelos, M., Pachitariu, M., 2021. Cellpose: a generalist algorithm for cellular segmentation. Nat. Methods 18, 100–106.

Tibshirani, R.J., Taylor, J., Lockhart, R., Tibshirani, R., 2014. Exact Post-Selection Inference for Sequential Regression Procedures. arXiv [stat.ME].

Wisniewski, J.R., Zougman, A., Nagaraj, N., Mann, M., 2009. Universal sample preparation method for proteome analysis. Nat. Methods 6, 359–362.

Yang, P., Humphrey, S.J., Cinghu, S., Pathania, R., Oldfield, A.J., Kumar, D., Perera, D., Yang, J.Y.H., James, D.E., Mann, M., Jothi, R., 2019a. Multi-omic Profiling Reveals Dynamics of the Phased Progression of Pluripotency. Cell Systems. https://doi.org/10.1016/j.cels.2019.03.012

Yang, P., Humphrey, S.J., Cinghu, S., Pathania, R., Oldfield, A.J., Kumar, D., Perera, D., Yang, J.Y.H., James, D.E., Mann, M., Jothi, R., 2019b. Multi-omic Profiling Reveals Dynamics of the Phased Progression of Pluripotency. Cell Syst 8, 427–445.e10.

Zarnegar, B.J., Flynn, R.A., Shen, Y., Do, B.T., Chang, H.Y., Khavari, P.A., 2016. irCLIP platform for efficient characterization of protein–RNA interactions. Nat. Methods 13, 489–492.

Zhu, A., Ibrahim, J.G., Love, M.I., 2019. Heavy-tailed prior distributions for sequence count data: removing the noise and preserving large differences. Bioinformatics 35, 2084–2092.

